# The exotic thymidine modification 5-hydroxymethyluridine in dinoflagellate *Amphidinium carterae*

**DOI:** 10.1101/2023.11.30.569493

**Authors:** Chongping Li, Ying Li, Yuci Wang, Ruixiang Meng, Xiaoyan Shi, Yangyi Zhang, Nan Liang, Hongda Huang, Yue Li, Hui Zhou, Jiawei Xu, Wenqi Xu, Hao Chen

**Author notes:** Corresponding author (C.P.L); (J.W.X.); (W.Q.X); (H.C.). These authors contributed equally to this work (Chongping Li, Ying Li, Yuci Wang, Ruixiang Meng).

## Abstract

Dinoflagellate chromosomes are extraordinary, as their organization is independent of architectural nucleosomes unlike typical eukaryotes and shows a cholesteric liquid crystal state. 5-hydroxymethyluridine (5hmU) is present at unusually high levels and its function remains an enigma in dinoflagellates chromosomal DNA. Here, we demonstrate that 5hmU exhibits content variations in different dinoflagellates and is generated at the poly-nucleotide level through hydroxylation of thymidine. Importantly, we identified the enzyme, which is a putative dinoflagellate TET/JBP homologue, catalyzing 5hmU production using either *in vivo* or *in vitro* biochemical assay. Based on the near-chromosomal level genome assembly of dinoflagellate *Amphidinium carterae*, we depicted a comprehensive 5hmU landscape and found that most 5hmU peaks share a conserved TG-rich motif, and are significantly enriched in repeat elements, which mark partially overlapping regions with 5-methylcytosine (5mC) sites. Moreover, inhibition of 5hmU via dioxygenase inhibitor leads to transcriptional activation of 5hmU-marked transposable elements (TEs), implying that 5hmU appears to serve as epigenetic marks for silencing retrotransposon. Together, our results revealed the biogenesis, genome-wide landscape and molecular function of dinoflagellate 5hmU, providing mechanic insight into the function of this enigmatic DNA mark.

## Introduction

In addition to the four canonical nucleobases including adenine, thymine, cytosine, and guanine, DNA molecules also contain different covalent modifications such as 5mC, N6-methyladenosine (6mA), *β*-D-glucosyl-hydroxymethyluracil (base J), 5-hydroxymethylcytosine (5hmC), 5-formylcytosine (5fC), 5-carboxylcytosine (5caC) and 5hmU in living organisms^1^. Methylation of cytosine to 5mC and adenine to 6mA are the most prevalent DNA modifications in eukaryotes and prokaryotes. 5mC is an ancient and conserved epigenetic mark that plays indispensable roles in the bacterial defense against phage attacks and eukaryotic transcription regulation ^2^. In contrast to the scarce amount of 6mA in animals and plants, 6mA is found extensively in bacteria and lower eukaryotes such as green algae and ciliates, and recognized as an important means to regulate gene expression, bacterial DNA repair/replication as well as eukaryotic nucleosome positioning ^3^. The non-canonical DNA modified base J in kinetoplastid protists also acts as an epigenetic mark to control variant surface glycoprotein (VSG) activation and RNA polymerase II (Pol II) transcription termination ^4,5^.

Due to the discordance between buoyant density and thermal denaturation temperature in the estimations of their genomic GC content, 5hmU was firstly found as a natural component of the DNA from bacteriophages, including SP8, SP01, Φe, H1, 2C and SP82, and responsible for the discrepancy ^6,7^. This conflict in estimating GC content was also observed in dinoflagellates and thus led to the discovery of 5hmU in their genomic DNA (gDNA) ^8–10^. It was reported that 5hmU could substitute for 12-68% of the thymidine in genomes of various dinoflagellates ^8–10^. In contrast, 5hmU content in other eukaryotes is much lower than in dinoflagellates, which only accounts for 500 - 7800 bases in mammalian genomes ^11^. As a precursor of base J, the average level of 5hmU is estimated to be about 0.02-0.12% of the total nucleotides in kinetoplastid flagellates *Trypanosomes brucei* ^12^. Consistent with the role as base J intermediates, genome-wide mapping of 5hmU loci in the parasitic kinetoplastid *Leishmania* demonstrated that they were mainly overlapped with base J-enriched loci, showing enrichment in stand switch/intergenic region and telomere ^13^. Despite the very high level of 5hmU, genome-wide mapping and functional investigation of 5hmU in dinoflagellates have not been described until now.

Previous studies showed that biosynthesis of 5hmU occurs through mechanisms operating either prior to or after DNA synthesis. In bacteriophages such as SP8 and SP01, they rely on utilization of intracellular dNTP pool to synthesize 5-hydroxymethyl-2’-deoxyuridine 5’-triphosphate (5hmUTP) ^14^. 5hmUTP production is initiated with deamination of deoxycytidine monophosphate (dCMP) to generate deoxyuridine monophosphate (dUMP), followed by dUMP hydroxymethylase activity to yield 5hmU monophosphate (5hmUMP) and then phosphorylation by 5hmUMP kinase. The resulting 5hmUTP is then incorporated into their genome during replication by a phage encoded DNA polymerase. Meanwhile, phage-related dTTP and dTMP nucleotidohydrolase (dTTPase and dTMPase) are activated to metabolize thymine nucleotides ^7^. On the other hand, 2-oxoglutarate/Fe(II)-dependent oxygenase (2-OG oxygenase) including JBP1 and JBP2 are responsible for conversion of thymidine into 5hmU at the DNA polymer level in kinetoplastids ^15^. Similarly, some phages can also utilize TET/JBP family proteins to install 5hmU in their chromatin after DNA synthesis^16^. However, the exact route of 5hmU generation in dinoflagellates remained an unsolved mystery.

Dinoflagellates, a major aquatic phytoplankton group with quasi-condensed liquid crystalline chromosomes (LCCs), are major primary producers in the oceans, essential photosymbionts of reef-building corals and also cause of harmful algal bloom. Dinoflagellate chromatin organization displays exceptional evolutionary divergence among eukaryotes. They behold some of the largest genome sizes, but are largely absent of architectural nucleosomes ^17^. With a low protein/chromatin DNA ratio of ∼1:10 and high levels of chromosomal divalent cations, dinoflagellate chromosomes occur in cholesteric liquid crystalline state, which are strongly birefringent and permanently condensed ^17,18^. Their genes are clustered into unidirectional arrays along the LCCs and lack of transcriptional regulation ^19–21^. Instead of the core histones, bacterial nucleoid-associated protein HU-derived “histone like proteins” (HLPs) and algae viruses-originated dinoflagellate viral nucleoproteins (DVNPs) constitute their major chromosomal packaging proteins ^17^. Additionally, dinoflagellates are the only eukaryotes with strikingly high amounts of DNA 5hmU modification, which could replace up to 68% of the thymidine (dT) in their LCCs ^17,18^. However, the contribution of 5hmU to the unique dinoflagellate LCCs and chromatin organization is still an enigma.

Since the initial discovery of the presence of 5hmU in dinoflagellate over 50 years ago, the encoding enzymes, genome-wide distribution pattern and role of this modification remained largely unexplored. In this study, we found that 5hmU also belongs to post-replicative enzymatic DNA modifications, which is catalyzed by the homologues of TET/JBP family proteins in dinoflagellates. Advances in next-generation sequencing have led to an explosion of available dinoflagellate genomes especially in *Symbiodinium* strains ^22^, which made the genomic-wide mapping of 5hmU possible. Here, based on long-read sequence assembly of dinoflagellate *A. carterae* genome, we found that 5hmU was preferentially deposited in intron as well as intergenic regions, and possibly served as an epigenetic mark to repress transposon activity. Our study thus provided the first integrated view of the biosynthesis, landscape and function regarding 5hmU in dinoflagellates.

## Results

### 5hmU is an abundant modification in Dinoflagellates genomic DNA

According to previous studies ^8,9^, besides the unusually high amounts of 5hmU in their nuclear DNA, dinoflagellates also contain non-neglectable levels of other DNA modifications. To further characterize the presence of diverse covalent modifications in dinoflagellates DNA, we first performed LC-MS/MS analyses to quantify the common DNA modification species in gDNA obtained from three different dinoflagellates including *A. carterae*, *Crypthecodinium cohnii* and *Symbiodinium* sp.. With the exception of 5fU and very low level of 6mA, the other DNA modifications, including 5hmU (Fig. 1A, 1B), 5mC and 5hmC, were readily detected in these organisms (Supplementary Fig. 1A, 1B, 1C, 1D, 1E). Being the most abundant DNA modifications, 5hmU accounts for 7.47%, 3.77% and 0.84% of their total thymidine (dT) in gDNA of *A. carterae*, *C. cohnii* and *Symbiodinium* sp. respectively (Fig. 1C), while 5mC constitutes 2.00%, 1.09%, and 1.05% of their total cytosine in these organisms (Fig. 1C). Additionally, we also found a relatively low amount of 5hmC (commonly less than 0.18% of total cytosine) was present in these three dinoflagellates (Supplementary Fig. 1E).

**Figure 1.**
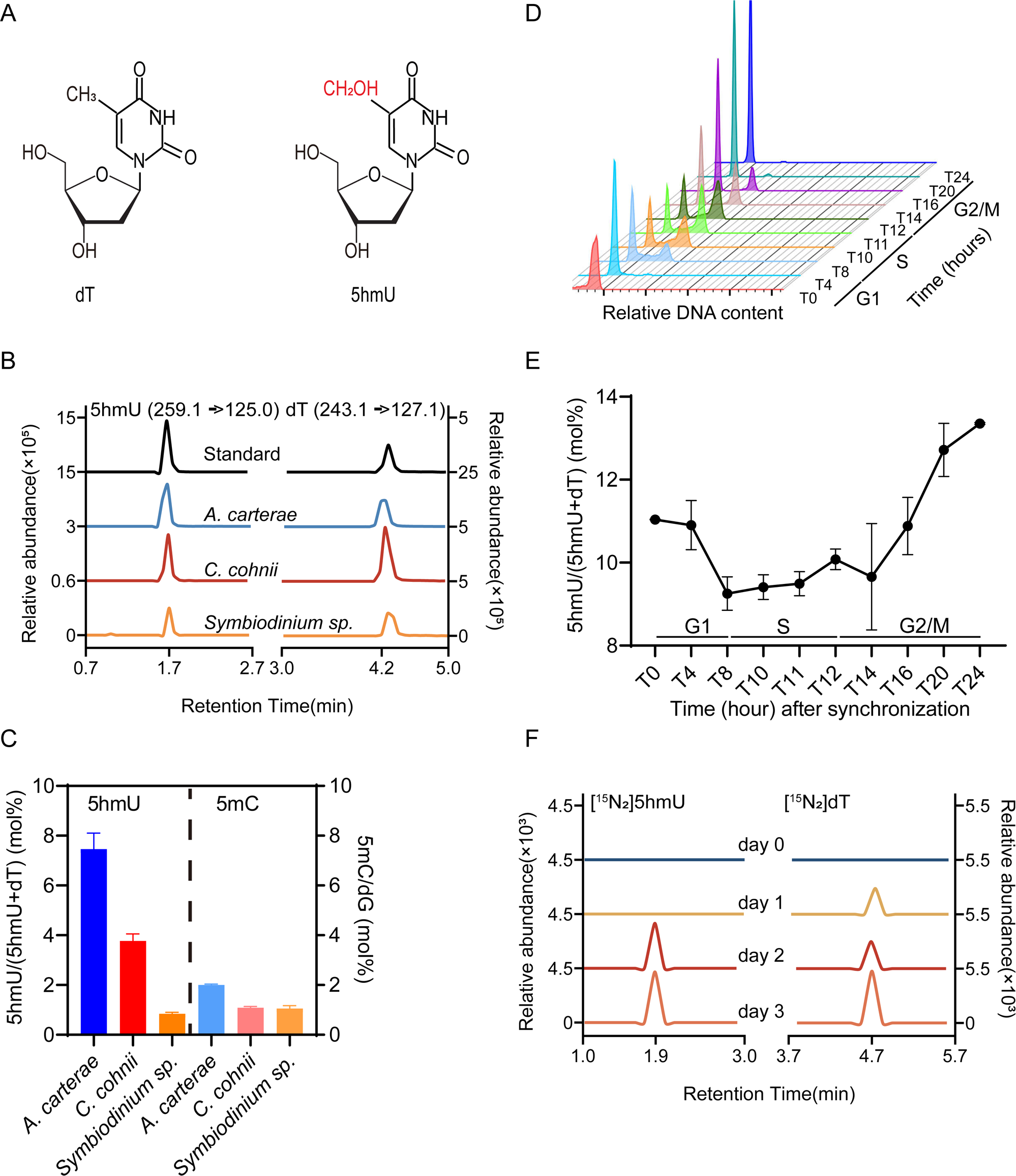
Characterization and quantification of 5hmU in Dinoflagellate gDNA. (A) Chemical structures of dT and 5hmU. (B) LC-MS/MS chromatograms of 5hmU and dT for different dinoflagellate species including *A. carterae*, *C. cohnii* and *Symbiodinium* sp.. (C) LC-MS/MS quantification analysis of 5hmU and 5mC in the gDNA of the mentioned three dinoflagellate species. Three biological replicates were performed. (D-E) DNA histograms and corresponding 5hmU content of the dinoflagellate *A. carterae* during the whole cell cycle. Two biological replicates were performed. (F) Detecting the presence of both heavy [^15^N_2_]5hmU and [^15^N_2_]dT in their isolated genomic DNA using LC-MS/MS after feeding *A. carterae* cells with nitrogen stable isotope-labeled thymidine ([^15^N_2_]dT).

To examine the dynamics of 5hmU during the cell growth, we measured the 5hmU level along the whole cell cycle of dinoflagellate *A. carterae.* Relative DNA content was firstly plotted against the time of cell cycle progression (Figure 1D). Flow-cytometric analyses showed that the DNA content of *A. carterae* cells kept stable at the G1 stage (T0 to T8), increased at the S phase (T8 to T12) and finally returned back to the original level during the G2/M phase (T12 to T24) (Figure 1D). LC-MS/MS quantification results indicated that 5hmU was preserved during the whole cell cycle and experienced a slight drop (about ∼16% decrease) of the 5hmU content at the corresponding S phase, followed by an increase of the 5hmU amount to its initial level at the G2/M stage (Figure 1E). This replication-coupled dilution and restoration of 5hmU in dinoflagellate *A. carterae* implied that 5hmU was established after new DNA synthesis and probably dependent on post-replicative enzymatic installation. Furthermore, it was reported that thymidine salvage pathway can enable cells to assimilate thymidine from the environment for dTTP synthesis ^23^. We determined to define the possible process of generating 5hmU through incubation of the *A. carterae* cells with nitrogen stable isotope-labeled thymidine ([^15^N_2_]dT). We found that with a 1-day interval, both [^15^N_2_]dT and [^15^N_2_]5hmU signals could be sequentially identified in their gDNA (Figure 1F, Supplementary Fig. 1F). This result further suggested that 5hmU was synthesized at the poly-nucleotide level through oxidation of thymidine.

### *De novo* genome assembly of dinoflagellate *A. carterae*

Among the three dinoflagellate species in this study, dinoflagellate *A. carterae* contains the highest 5hmU content, but no reference genome is available for now. In order to further characterize the genome-wide profile of 5hmU, we sought to decode the whole genome sequence of *A. carterae*. The genome size was firstly determined using flow cytometry and estimated to be about 2.1 Gbp by nominalization against human iPSC (Induced pluripotent stem cell) DNA content (Supplementary Fig. 2A). Genome survey with k-mer counts analysis based on the short read-sequencing data indicated that *A. carterae* was haploid and had a genome size about 1.21 Gbp (Supplementary Fig. 2B). This genome size estimation discordance was also observed in other dinoflagellates and possibly attributed to the liquid crystal structure of dinoflagellate chromosomes ^24^.

A nearly chromosome-scale genome of *A. carterae* was obtained by combing short- and long-read sequencing, and Hi-C-assisted assembly (Fig. 2A). The final *A. carterae* assembly is about 1.26 Gbp (mean GC content = 44.81%, Fig. 2B) and contains a total of 314 scaffolds (scaffold N50 = 34.285 Mb and contig N50 = 820.60 Kb; Figure 2C, Supplementary Fig. 2C), of which 43 were identified as major pseudo-chromosomes comprising ∼98.17% of the total genome sequence (Fig. 2A, Supplementary Fig. 2C). This assembled genome is highly repetitive and consists of about 56% repeat sequences, which mainly include long terminal repeat (LTR) elements, such as Copia and Gypsy retrotransposons, long interspersed nuclear elements (LINE), DNA transposons (DNA TE) and unknown repeats (Fig. 2C). Gene predication yielded 41274 protein-coding genes (Supplementary Fig. 2C), of which 79.83% had close matches in reference databases. Additionally, these genes showed a higher average GC content (∼54.56%) than that of the whole genome sequence (Fig. 2B). Consistent with other dinoflagellates ^24,25^, most of these genes use GC in addition to GT as the 5’ splice donor motifs (Fig. 2D) and are unidirectionally encoded, exhibiting seldom gene orientation change (Fig. 2E) and approximately equal distribution of genes in both gDNA strands (Fig. 2F).

**Figure 2.**
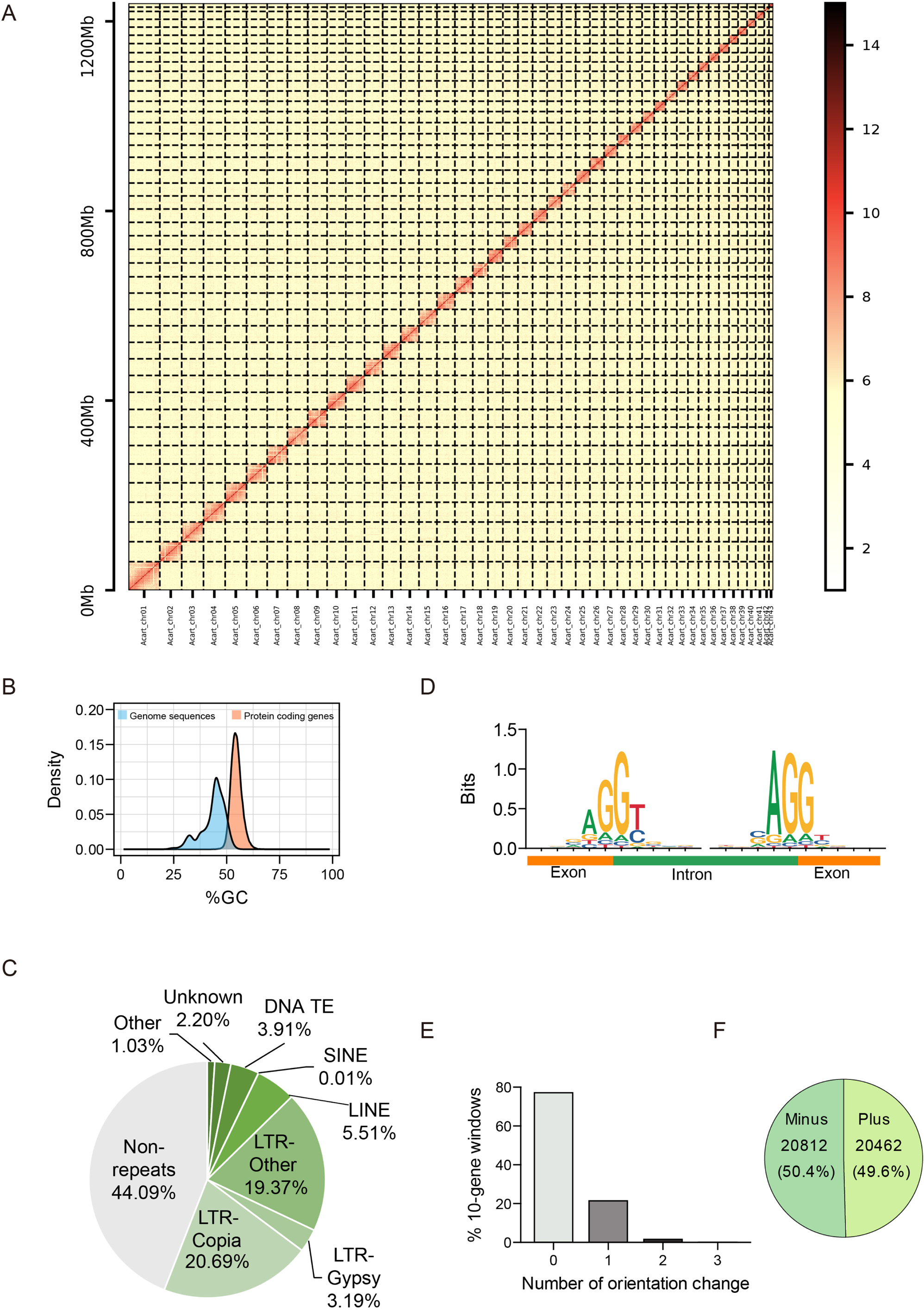
*De novo* assembly and characteristics of dinoflagellate *A. carterae* genome. (A) Hi-C interaction map of 43 chromosome-scale scaffolds (ordered by descending size, 500-kb bins) for *A. carterae.* (B) Average GC content distribution of *A. carterae* genome sequence for each pseudochromosomes and their protein coding genes respectively. (C) Proportion of non-repeat sequence and different repeat classes in the assembled genome of *A. carterae*. (D) The consensus sequences of the splice sites in all annotated genes of *A. carterae*. (E) Frequency of gene orientation changes in 10-gene windows across the *A. carterae* genome. (F) Pie chart representation of gene directionality.

### Putative dinoflagellate Tet/JBP family protein homologues are the thymidine hydroxylases involved in 5hmU biosynthesis

It has long been known that TET/JBP family proteins have enzyme activities toward thymidine in the context of DNA polymer ^26^. To identify the enzymes responsible for the addition of 5hmU in dinoflagellates, we first examined the presence of Tet/JBP family proteins in dinoflagellates genomes. Within dinoflagellates *C. cohnii* and *A. carterae* used in this study, iterative sequence similarity search using the oxygenase domain of TET/JBP superfamily recovered homologous regions in two proteins, which were named as *Crypthecodinium cohnii* TET/JBP homologue 1 (CryTJ1) and *Amphidinium carterae* homolog 1 (AmpTJ1), respectively ( Fig. 3A), Their numerous orthologues were also identified across different sister taxa (Supplementary Fig. 3A), which formed several distinct clades during the evolution (Supplementary Fig. 3A). Multiple sequence alignment also revealed the conserved signature motifs including HXD and H•••R for binding cofactors Fe^2+^ and 2-oxoglutarate (2OG) in these two newly predicted TET/JBP homologues CryTJ1 and AmpTJ1 (Supplementary Fig. 3B). Thus, despite the highly divergent sequence with typical eukaryotes, we successfully identified a variety of dinoflagellate TET/JBP homologues.

**Figure 3.**
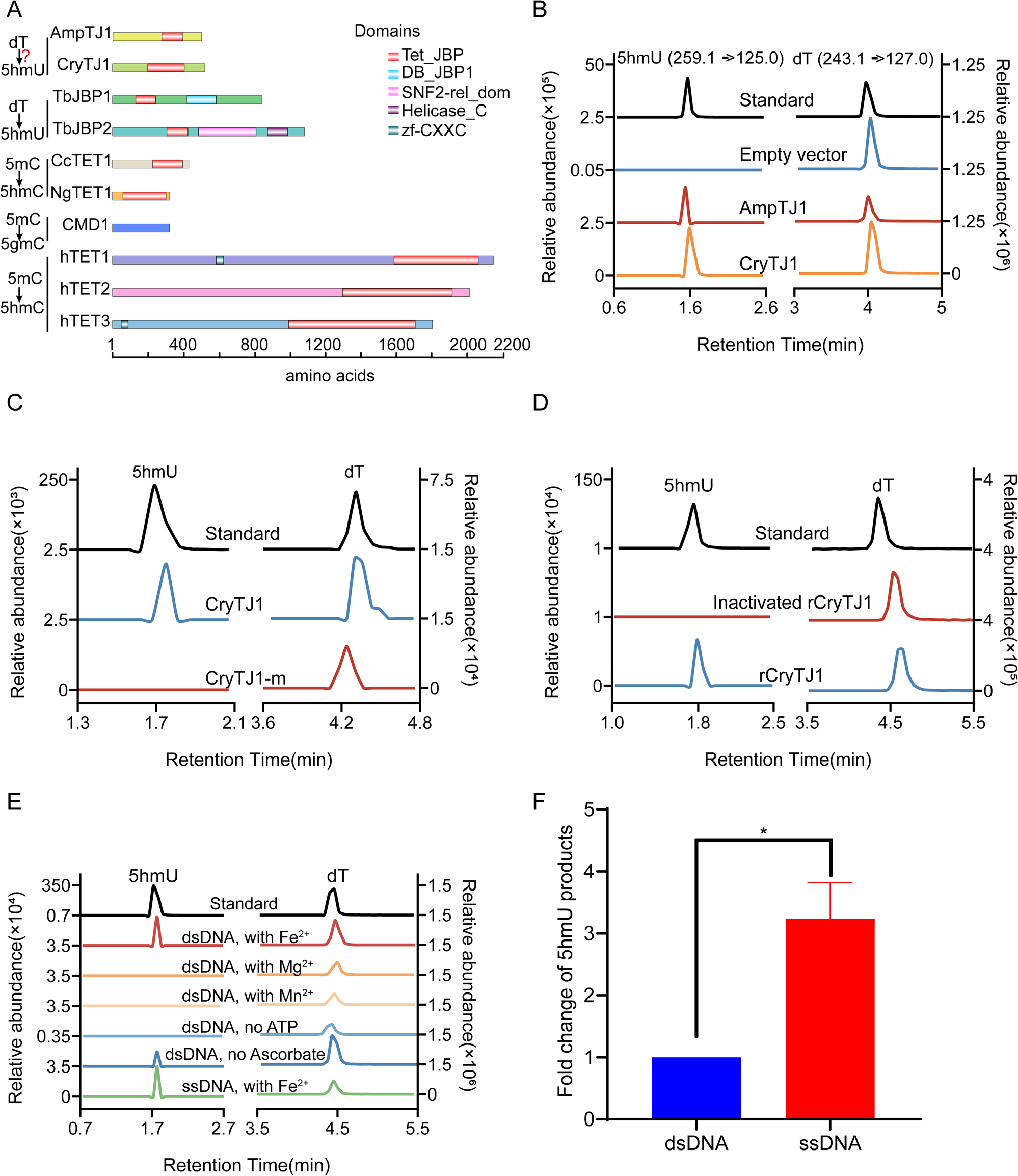
Characterization of the thymidine hydroxylation activities of dinoflagellate Tet/JBP homologues AmpTJ1 and CryTJ1. (A) Schematic representation of the domain architecture of AmpTJ, CryTJ1 and other representative Tet/JBP family proteins with known enzyme activities toward thymidine/5mC including TbJBP1 (P86937) and JBP2 (Q57X81) from *Trypanosoma brucei*, ccTET1 (XP001838108.2) from *Coprinopsis cinerea*, NgTET1 (XP002667965.1) from *Naegleria gruberi*, CMD1 (Cre12.g553400.t2.1) from *Chlamydomonas reinhardtii*, and hTET1 (Q8NFU7), hTET2 (Q6N021) as well as hTET3 (O43151) from *Homo sapiens*. 5gmC, abbreviations of 5-glyceryl-methylcytosine. (B) LC-MS/MS analysis of the *in vivo* thymidine oxidation activity for *E. coli* strains containing empty vector, pET28a-AmpTJ1 and pET28a-CryTJ1 after IPTG induction. (C) A comparison of their thymidine hydroxylation activity between CryTJ1 and CryTJ1m carrying HDR to ARA mutation after IPTG induction of the corresponding recombinant *E. coli* strains. (D) *In vitro* biochemical assay of the purified rCryTJ1 using HEK293T gDNA as substrate. The heat inactivated form of rCryTJ1 was used as control. (E) The effects of different metal ions, the absence of ATP or ascorbate, and ssDNA substrate on in vitro enzyme activities of rCryTJ1. Fe^2+^, Mg^2+^, and Mn^2+^ represent Fe(NH_4_)_2_(SO_4_)_2_, MgCl_2_ and MnCl_2_ respectively. (F) Bar graph showing the differential enzyme activities of rCryTJ1 on dsDNA and ssDNA substrates (with paired t-test, p* < 0.05). Three biological replicates were performed.

To confirm the ability of dinoflagellate TET/JBP homologues to catalyze thymidine hydroxylation at DNA polymer level, CryTJ1 and AmpTJ1 were individually expressed in *Escherichia coli*, of which gDNA were extracted and used to validate 5hmU generation. LC-MS/MS analyses showed that both proteins can result in detectable thymidine hydromethylation of the gDNA with relatively higher enzyme activity of CryTJ1 than AmpTJ1 (Fig. 3B). Meanwhile, *E. coli* cells expressing CryTJ1 mutant (CryTJ1-m, bearing ARA residues rather than critical HRD motif in CryTJ1) abolished its capacity to synthesize new 5hmU (Fig. 3C). On the other hand, full length of recombinant CryTJ1 proteins (rCryTJ1) were purified with affinity chromatography (Supplementary Fig. 3C) and in *vitro* biochemical assay substantiated its capability for formation of 5hmU but not 5hmC with HEK293T gDNA as substrate (Fig. 3D, Supplementary Fig. 3D). Next, we compared the enzyme activity of rCryTJ1 with different metal ions and found that Fe^2+^ was essential for 5hmU production (Fig. 3E). Meanwhile, the catalytic activity was also negatively affected by the absence of ATP or cofactor ascorbate in the reaction system (Fig. 3E). Interestingly, compared with double-stranded DNA (dsDNA), rCryTJ1 exhibited significantly higher enzyme activity towards single-stranded DNA (ssDNA) substrate, indicating a preference for ssDNA structure (Fig. 3F). This discovery is consistent with previous structural and biochemical studies concerning other TET/JBP family members, which contact with 5mC on only one strand when the catalytic reaction occurs ^27,28^. Taken together, these results demonstrated that dinoflagellate TET/JBP homologues could convert thymidine to 5hmU at the polynucleotide level through oxidation and probably served as the enzymes responsible for dinoflagellate 5hmU production.

To explore whether CryTJ1 could carry out intracellular 5hmU synthesis on nucleosome-wrapped chromosomes, we constructed a stable HEK293T cell line containing a Doxycycline (Dox)-inducible Flag-tagged CryTJ1 with three SV40 nuclear localization signals (3×NLS-CryTJ1-Flag), and the expected protein bands were confirmed with immunoblot following Dox induction, but not the empty vector control (Fig. 4A). Immunofluorescence analysis also indicated the successful localization of CryTJ1 within the nuclei of HEK293T cells (Fig. 4B). However, similar with the empty vector and non-doxycycline induction control groups, no detectable 5hmU could be detected in stable HEK293T cell line expressing Flag-tagged CryTJ1 (Fig. 4C), indicating nucleosomes seemed to be obstacles for 5hmU production. Meanwhile, we transformed budding yeast cells with a green fluorescent protein (GFP) fused version of 3×NLS-CryTJ1-Flag (3×NLS-CryTJ1-GFP-Flag). With successful expression and co-localization with the yeast nuclei (Supplementary Fig. 4A, Fig. 4B), the CryTJ1 also failed to install 5hmU in gDNA of these budding yeast cells (Supplementary Fig. 4C). Consistently, *in vitro* biochemical assay using the naked DNA, but not mono-nucleosome, as substrate led to 5hmU synthesis (Fig. 4D). Thus, our results suggested that nucleosome-protected DNA is not the preferred substrate of the 5hmU synthase, which may be related to the adaption of this enzyme’s activity against the nucleosome-less LCCs in dinoflagellates.

**Figure 4.**
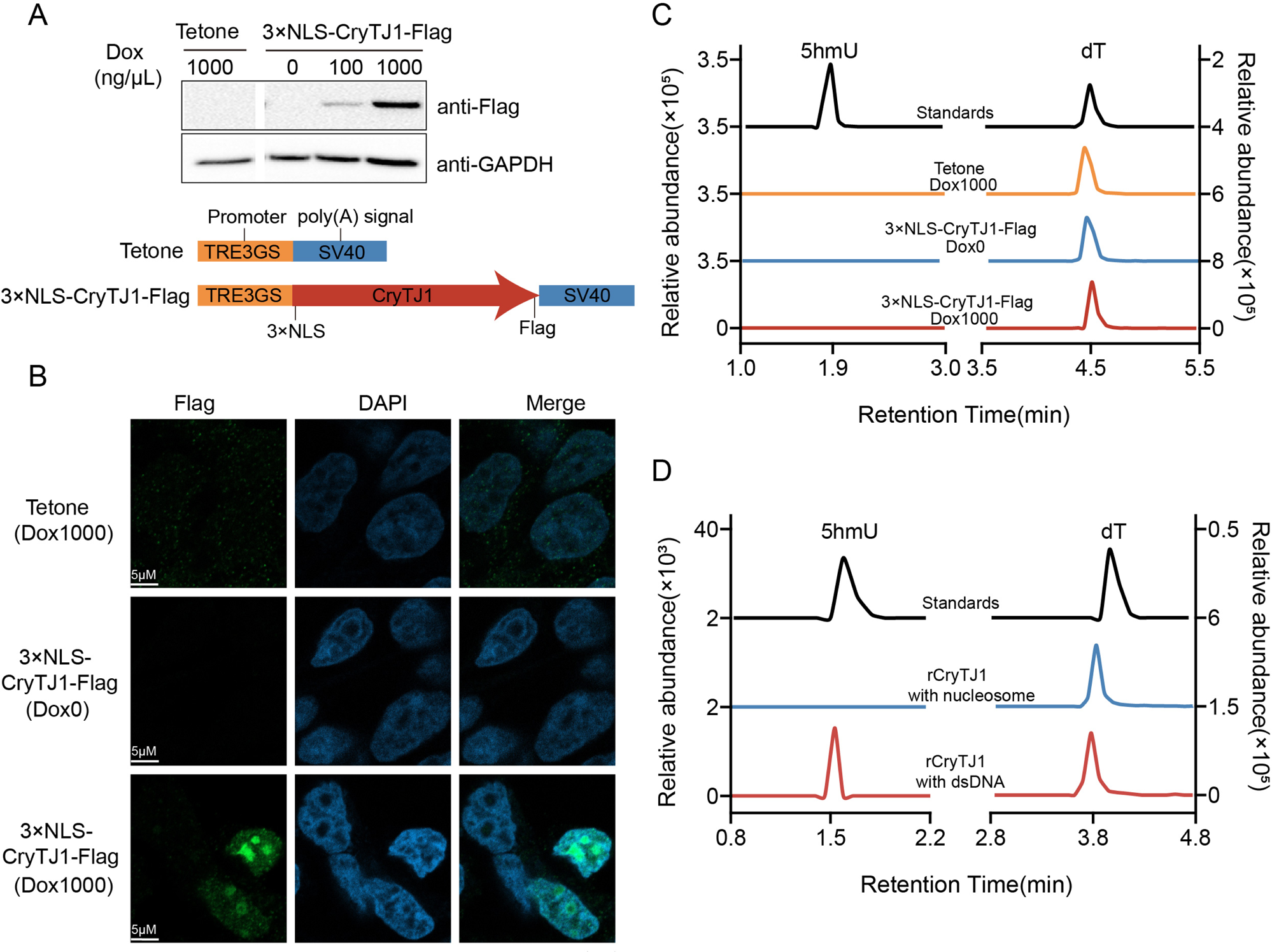
Wrapped nucleosomal DNA hinders the formation of 5hmU by CryTJ1. (A) Western blot analysis of HEK293T cells expressing either empty vector pLVX-Tetone-Puro (Tetone) or Flag-tagged CryTJ1 with three SV40 nuclear localization signals (3×NLS-CryTJ1-Flag) after doxycycline (dox) induction. GAPDH was used as loading controls. Schematic representation of the expression cassettes with the TRE3GS promoter and SV40 poly(A) signal sequences for plasmids Tetone and pLVX-3×NLS-CryTJ1-Flag-Tetone-Puro were shown below the immunoblots. (B) Immunofluorescence micrographs showing colocalization between the Flag epitopes on CryTJ1 (green) and DAPI stain (blue). (C) LC-MS/MS chromatograms displaying the no appearance of 5hmU after dox induction for different pLVX-Tetone-puro constructs. (D) A comparison of the enzymatic activities of purified rCryTJ1 proteins on substrates including mononucleosome and the corresponding naked dsDNA.

### Genome-wide mapping of 5hmU in dinoflagellate *A. carterae*

Although 5hmU is highly abundant in dinoflagellates, the exact distribution pattern remains obscure, which hinders the illustration of 5hmU’s regulatory roles. With similar pyrimidine structures and the ability to fully oxidize dsDNA 5hmC sites into 5fC by mild chemical 4-acetamido-2,2,6,6-tetramethylpiperidine-1-oxoammonium tetrafluoroborate (ACT^+^BF4^−^) ^29^, we envisaged this chemical could also be utilized to carry out complete conversion of *A. carterae* gDNA 5hmU into 5fU without denaturation. Subsequent *in vitro* oxidation assay successfully confirmed that chemical ACT^+^BF4^−^ were able to fully oxidize 5hmU into 5fU at the dsDNA polymer level (Supplementary Fig. 5A). To map the genome-wide 5hmU distribution, we utilized an anti-5hmU antibody-based DNA 5hmU immunoprecipitation (5hmU-DIP) method and set up a parallel DIP control with 5fU-converted gDNA (named as 5fU-DIP group) to unveil the 5hmU landscape and mask the non-specificity of 5hmU antibody (Supplementary Fig. 5B). After comparing both the 5hmU-DIP and 5fU-DIP reads with the sequencing background (input) individually, we found 5hmU-DIP captured highly specific signals, which was largely different from 5fU-DIP signals (Supplementary Fig. 5C). The 5hmU-DIP analysis results indicated that more than 97% of 35642 5hmU peaks were enriched in intergenic region and intron (Fig. 5A, 5B), implying a non-random distribution pattern of 5hmU along the *A. carterae* chromosomes. It was also found that there was a highly conserved motif GTTGTTGTTG in these 5hmU-specific loci (Fig. 5C). Despite the remarkable difference in their total genomic 5hmU contents, this preference of G and T bases resembles the 5hmU motif signatures observed in the evolutionally distant kinetoplastid *Leishmania major* ^30^.

**Figure 5.**
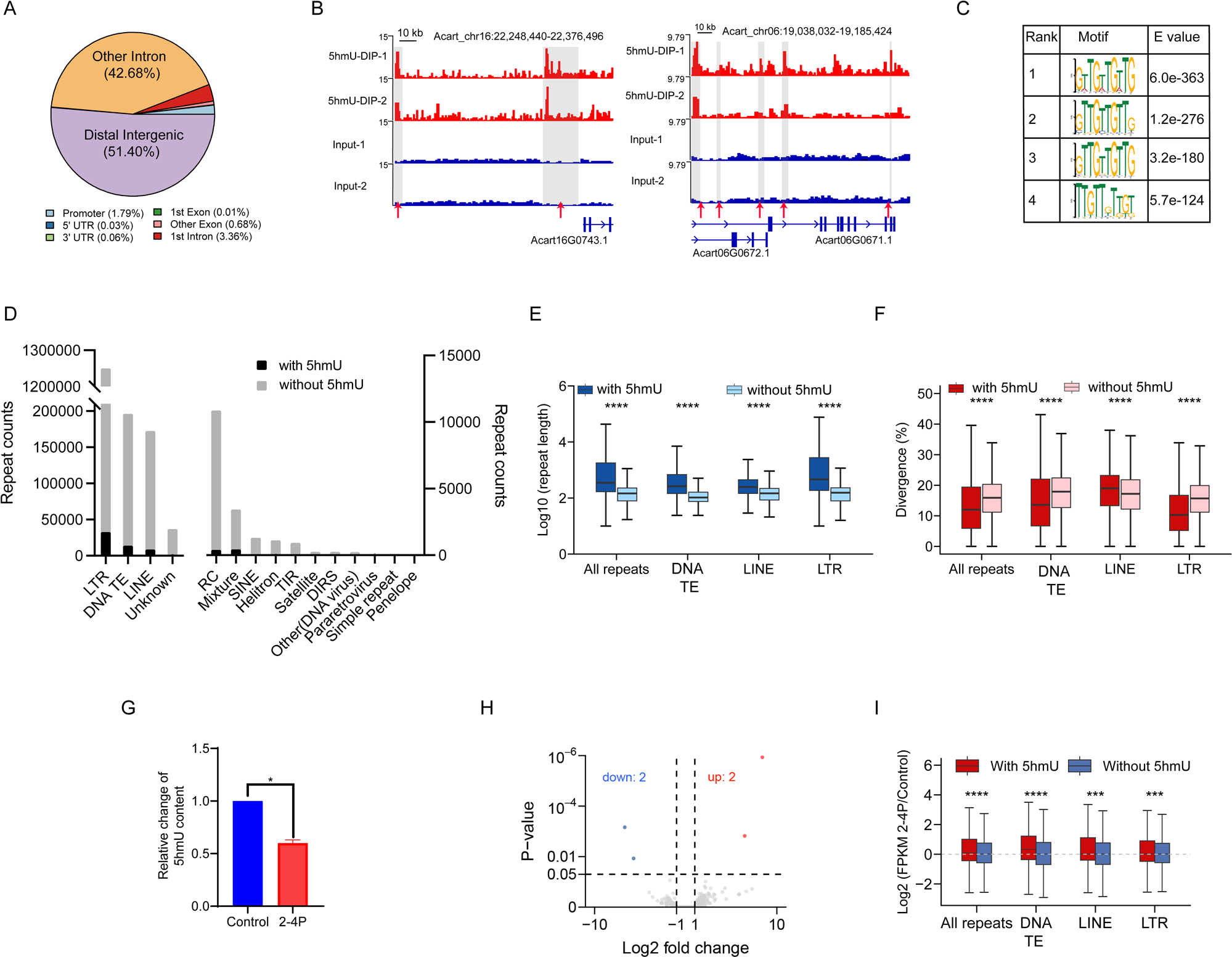
Genomic landscape of 5hmU in dinoflagellate *A. carterae*. (A) Distribution of 5hmU peaks across different genomic regions. (B) Representative 5hmU-DIP profiles in selected genomic regions. The 5hmU peaks were marked by read arrows. Two biological replicates were shown for both 5hmU-DIP and input samples respectively. (C) Conserved motifs for genome-wide 5hmU peaks. (D) The distribution pattern of distinct repeat classes with or without 5hmU peaks. (E and F) The length (E), and corresponding divergence (F) of all repeats, DNA transposon, LINE and LTR with or without 5hmU peaks (Wilcox test, ****p < 0.0001). (G) The decrease of global 5hmU levels after 2-4P treatment (with paired t-test, p* < 0.05). Control, *A. carterae* cells with no 2-4P treatment. 2-4P, *A. carterae* cells with 2-4P treatment. Two biological replicates were performed. (H) Volcano plot of the distribution of differential expressed genes, mapping the 2 upregulated genes (red color) and 2 downregulated genes (blue color). *A. carterae* cells treated with 2-4P vs untreated *A. carterae* cells as Controls. (I) The Log2 fold change of transcriptional expression levels (FPKM values) for *A. carterae* cells after 2-4P treatment regarding all repeats, DNA transposon, LINE and LTR with or without 5hmU peaks (Wilcox test, ***p < 0.001 and ****p < 0.0001). For all boxplots, the center line is the median. The boxes upper and bottom boundaries represent the first quartile and third quartile, while the error bars denote the highest and lowest values.

Meanwhile, we also carried out an independent chemical-tagging strategy to identify the genome-wide 5hmU loci in *A. carterae* (named as 5hmU-chemical-seq, Supplementary Fig. 6A). Though we only found a total of 2892 5hmU high-confidence peaks using this method, the recovered motifs (TG-rich) remained almost identical and 1311 peaks shared overlapped position with 5hmU-DIP peaks (Supplementary Fig. 6B, 6C). Moreover, a 5hmU-specific loci mapping based on 5hmU-DIP approach was also performed in the previous *Symbiodinium* sp. strains used in this study, and the results showed that although only 35.43%-44.38% of clean reads could be mapped to the reference genome of *Effrenium voratum* CCMP421 ^31^, a total of 2183 5hmU peaks were recaptured, sharing similar motifs to those of *A. carterae*, of which the majority were also enriched within intergenic and intronic regions (Supplementary Fig. 7A, 7B). Thus, significantly fewer 5hmU peaks are determined in *Symbiodinium* sp., which is consistent with the LC-MS/MS result that *Symbiodinium* contains a significantly lower level of 5hmU than *A. carterae* (Fig. 1C). Cumulatively, the results indicated that dinoflagellate 5hmU have preferential localization in chromosomes and evolutionarily conserved TG motifs, which was possibly attributed to the similar substrate requirement of TET/JBP family proteins.

Given the preferential enrichment of 5hmU peaks in intergenic and intronic regions, we speculated whether 5hmU overlapped with the repetitive regions in the assembled genome. Interestingly, we found that 85.8% of the 5hmU peaks were distributed within the repeat sequences (Supplementary Fig. 8A), suggesting a major co-localization with the repeat elements. Specifically, 5hmU peaks showed preferential enrichment in TEs including retrotransposon LTR, DNA TEs and LINE elements (Fig. 5D). A comparative analysis between the repeat elements with or without 5hmU sites indicated that except the elevated divergence of LINE elements with 5hmU peaks, the repeat elements containing 5hmU marks normally have longer length and less divergence, which implied that the 5hmU-marked TEs were inserted more recently and likely still have activities (Fig. 5E, 5F, Supplementary Fig. 8B, 8C).

To further characterize the influence of 5hmU on repeat elements, we used a 5hmU synthase inhibitor 2,4-pyridinedicarboxylic acid hydrate (2-4P), to suppress the deposition of 5hmU in *A. carterae* genomic DNA (Fig. 5G). Then, RNA-seq analyses demonstrated that 5hmU content reduction didn’t cause significant transcriptional change for the protein coding genes, in which only a total of 4 genes showed differential gene expression (Fig. 5H). However, we observed a globally increased expression for the 5hmU-decorated TEs (Fig. 5I). Specifically, DNA TE, LTR retrotransposons such as Copia and Gypsy (Supplementary Fig. 8D), and LINE showed elevation at their transcriptional level after 2-4P treatment (Fig. 5I), implicating 5hmU may act as an epigenetic mark to repress their transcriptional activity.

### Correlation between 5hmU and 5mC DNA marks

Given the non-ignored amount of 5mC present in *A. carterae* genome, we characterized the DNA 5mC methylomes and examined whether there is any connection between these two DNA marks. With methylation detected in three cytosine contexts, the CG, CHG, and CHH methylation occurred at the highest, medium and lowest levels in *A. carterae* respectively (Supplementary Fig. 9A). Their CG methylation (mCG, ∼64.08%) was close to the methylation level reported in dinoflagellate *S. minutum* ^32^, and also showed a unimodal distribution (Supplementary Fig. 9A, 9B). In contrast, methylation in both CHG (mCHG) and CHH (mCHH) reached 6.26% and 3.89% respectively (Supplementary Fig. 9A), which were higher than the corresponding levels of *Symbiodinium* strains (∼1%) ^32^. Specifically, the majority of CG sites exhibited medium to high levels of cytosine methylation, while only about 0.57% of CGs being non-methylated (Supplementary Fig. 9B), indicating a default methylation state of these cytosine sites. Additionally, mCG was evenly distributed along the genome and exhibited no association with gene expression (Supplementary Fig. 9C, 9E). However, 5mC in both CHG and CHH contexts showed significant enrichment in the TEs (Supplementary Fig. 9D), and their high methylation levels tended to inhibit transcription of corresponding TEs (Supplementary Fig. 9E). Consistent with the previous study ^32^, these results suggested that despite the accumulation of high levels of methylation at CG sites along the *A. carterae* genome, only the methylation levels of infrequent mCHG and mCHH were likely associated with transposon transcription.

Furthermore, a comparison of 5mC sites and 5hmU loci in genome-wide TEs of *A. carterae* suggested that 5mC marked repeat elements can be also decorated by 5hmU (Fig. 6A, 6B, 6C; Supplementary Fig. 10A, 10B, 10C, 10D). Generally, the TEs, either all repeats or DNA TE, LINE, Copia and Gypsy, containing both mCG sites and 5hmU peaks showed slightly reduced methylation level than these with exclusive mCG sites (Fig.6A; Supplementary Fig. 10A). On the contrary, the same set of TEs consisting of colocalized mCH (mCHG and mCHH) sites and 5hmU loci displayed increased methylation level when compared with the other TEs bearing only mCH marks (Fig. 6B, 6C; Supplementary Fig. 10B, 10C). Notably, 5hmU signals remained at high levels in the internal regions of repeat elements including DNA TE, LINE, Copia and Gypsy (Fig.6D). Specifically, both 5hmU and mCH signals are enriched at the internal regions of DNA TE and Gypsy elements, while 5hmU signals seemed to exhibit a near-complementary profile to the relatively lower levels of mCH in the internal regions of both LINE and Copia elements (Fig.6D). Moreover, 5hmU colocalization preferred to occur on both LINE and Copia elements with longer length (Fig.6E, 6F). Of note, except mCHH sites, LINE carrying with co-localized 5mC sites and 5hmU peaks tended to have higher divergence (Fig.6G), while Copia elements bearing overlapped 5mC sites and 5hmU peaks have less divergence (Fig.6H), which resembles the described features of corresponding 5hmU-containing TEs shown in previous figures (Fig. 5E, 5F). Thus, these results implied that 5hmU modification may cooperate with cytosine methylation mCH (but not mCG) together to regulate TE activity.

**Figure 6.**
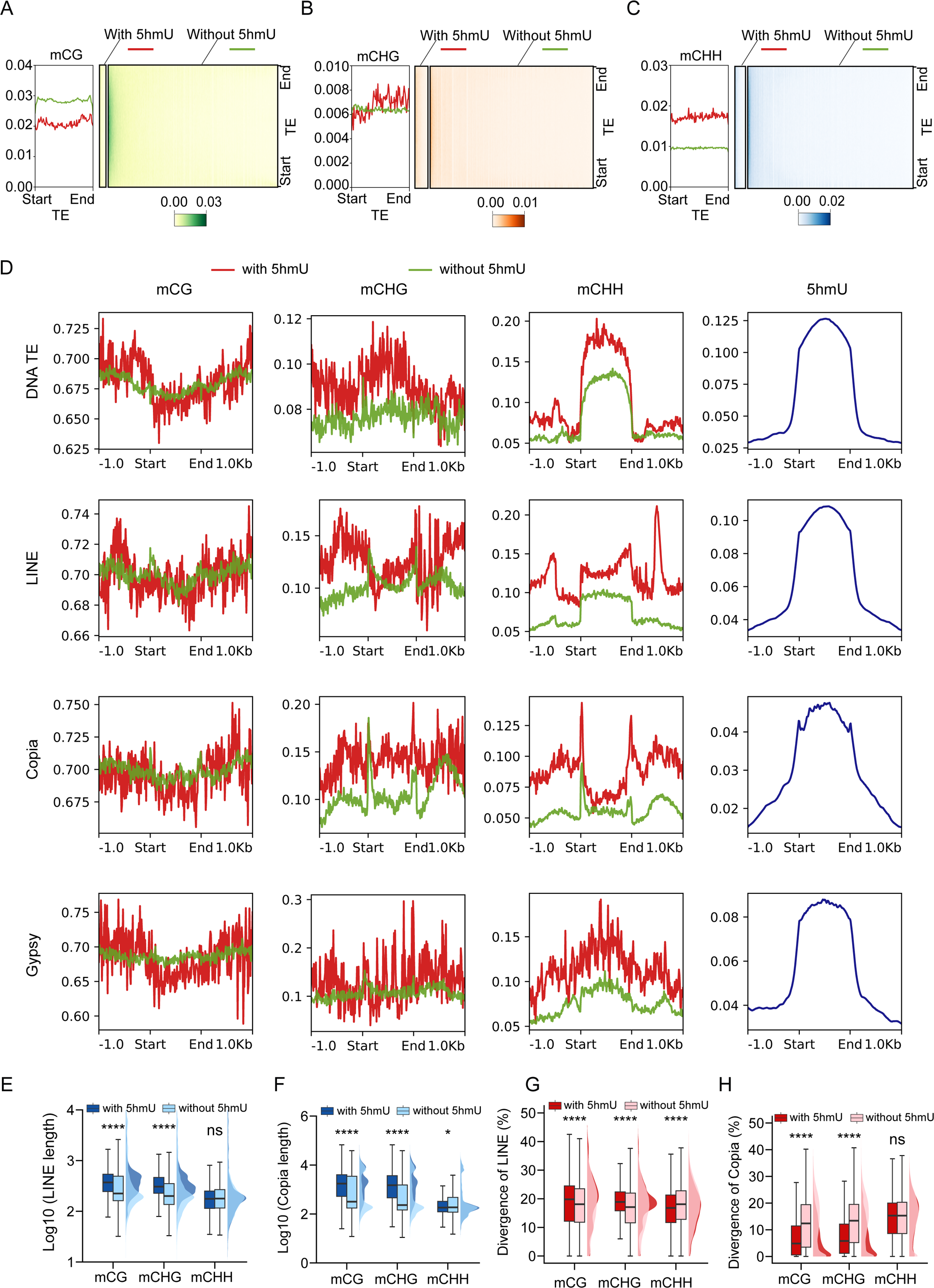
Comparison of genome-wide 5hmU and 5mC in dinoflagellate *A. carterae*. (A-C) Average density plot and heatmap of cytosine methylation profiles at CG (A), CHG (B) and CHH (C) sites in transposable elements (TEs) with or without 5hmU peaks. (D) Metaplots of 5hmU enrichment over different types of TEs compared to mCG, mCHG and mCHH sites. (E and F) Box plots showing the length of LINE (E) and Copia elements (F) with or without 5hmU peaks (Wilcox test, ns = not significant; *p <0.05; ****p <0.0001). (G and H) Box plots showing the corresponding divergence of LINE (G) and Copia elements (H) with or without 5hmU peaks (Wilcox test, ns = not significant; ****p <0.0001). For all boxplots, the centerlines are medians, box limits are quartiles 1 and 3, whiskers are the 1.5× interquartile and the outliers are not shown.

## Discussion

As a natural feature of dinoflagellates, the DNA of many dinoflagellates contains strikingly high amounts of 5hmU ^8,9^. While earlier measurements of 5hmU were primarily dependent on semi-quantitative methods (e.g., thin layer chromatography) ^8,9^, we substantiated the prevalent presence of 5hmU, and also found the existence of 5mC, 5hmC and 6mA, but not 5fU with ultra-sensitive LC-MS/MS analyses in three representative dinoflagellates. It was reported that about 68% and 37% of thymidine are replaced by 5hmU in dinoflagellates *A. carterae* and *C. cohnii* respectively, which is about 10-times higher than the corresponding 5hmU levels in the present study ^9^. This difference might be due to different strains and different methods for 5hmU detection used in these studies. 5hmU in living organisms is produced at either the mononucleotide level ^14^ based on the *de novo* synthesis of 5hmUTP in some bacteriophages or the polynucleotide level with the utilization of TET/JBP family proteins ^26^. The slight 5hmU level drop during S phase, the detection of heavy [^15^N_2_]5hmU sites after DNA incorporation of [^15^N_2_]dT, as well as the presence of functional TET/JBP homologues in dinoflagellates *A. carterae* and *C. cohnii* all point to the fact that dinoflagellate 5hmU is enzymatically synthesized through thymidine oxidation at the post-replication stage (Supplementary Fig. 11).

TET/JBP proteins were known to catalyze oxidative modification of 5mC or thymidine in chromosomal DNA ^26,33^. Although with highly divergent sequences, we successfully identified various TET/JBP homologues in dinoflagellates and confirmed the thymidine hydroxylase activities of CryTJ1 and AmpTJ1 proteins (Supplementary Fig. 11). For many epigenetic modifications, there are complicated regulatory networks involved in their *de novo* generation and maintenance. As to 5mC methylation, it’s well known that DNA methyltransferases DNMT3 and DNMT1 are responsible for 5mC establishment and maintenance respectively ^34^. Similarly, in kinetoplastid *Trypanosoma brucei*, there are two JBP proteins involved in synthesis of 5hmU, of which JBP2 is responsible for the *de novo* 5hmU and base J synthesis, while JBP1 only hydroxylates J-adjacent thymidine residues to generate 5hmU as the precursor for base J ^15^. There are also other different TET/JBP proteins in dinoflagellates, but the enzymatic activity towards thymidine and division of labor between these TET/JBP proteins in dinoflagellates await further investigation. It should be noted that dinoflagellates also contain considerable levels of 5hmC in the genome, and whether certain TET/JBP family member(s) act as 5mC-specific hydroxylase remains to be explored. Dinoflagellate LCCs contain barely detectable architectural nucleosomes, but low amounts of nucleoproteins such as HLPs and DVNPs ^17^. As CryTJ1 prefers to use non-nucleosomal DNA as its substrate, it would be interesting to further examine if the hydroxylase activity of dinoflagellate TET/JBP homologues can be affected by their chromosomal packaging proteins.

Previous studies in dinoflagellate *C. cohnii* showed that 5hmU was not randomly distributed, with variation of its amounts in different buoyant density fractions of isolated DNA and preferential replacement of thymine in dinucleotides TA and TC ^8,35^. Though 5hmU is suggested to be the remnant of an ancient restriction-modification system and has little or no contemporary function in dinoflagellates ^36^, our current study clearly argued against this speculation. In this study, we demonstrated that most 5hmU peaks were selectively enriched at intergenic and intronic regions of *A. carterae* genome and exhibited a conserved TG-rich motif (Supplementary Fig. 11), which was also observed in another dinoflagellate-*Symbiodinium* sp.. Thus, this non-random distribution pattern of 5hmU seemed to be conserved and likely of functional relevance in different dinoflagellates. The dinoflagellates commonly have giant genome size and it was hypothesized that repeat expansion contributes much to the genome size expansion ^22^. With main co-localization between 5hmU loci and repeat elements in *A. carterae* genome, we found that a ∼50% reduction of 5hmU content resulted in little variations in gene expression, but elevated expression of the TEs globally, suggesting 5hmU may work as an epigenetic mark to repress transposon activity at the transcriptional level (Supplementary Fig. 11). Interestingly, it has been shown that 5hmU is indeed involved in transcription regulation. An *in vitro* transcription assay with bacterial RNA polymerase indicated that DNA templates containing 5hmU could either increase or inhibit transcription in a context-dependent manner ^37^. However, it remains to be determined how 5hmU negatively affects the expression of TEs in dinoflagellates.

With CG hypermethylation as a default state and the homogenous distribution of mCG along the genome, dinoflagellate CG methylation appears to be not associated with either gene expression levels or transposon activation, which is unlike the typical function of mCG in other eukaryotes. While both mCHG and mCHH methylation in dinoflagellate *A. carterae* are negatively correlated with transposon transcriptional levels. We found that 5mC and 5hmU occupy overlapped repeat elements, implicating complementary roles of both modifications, especially the interaction of 5hmU and 5mC marks within both CHG and CHH contexts, in regulating transcription activities of TEs (Supplementary Fig. 11). 5hmU modifications were known to increase flexibility and hydrophilicity of dsDNA structure ^38^. In cooperation with high concentrations of chromosomal divalent cations, 5hmU was believed to promote DNA twistability of dinoflagellate LCCs ^18^. TEs play important roles in maintaining chromosomal organization including demarcating topologically associating domain (TAD) boundaries, promoting short or long-range interactions and maintaining heterochromatin stability ^39^. With a dip of GC content, the convergent strand switch regions of unidirectional gene cluster delimit dinoflagellate TADs boundaries ^19,20^. Therefore, how these 5hmU modifications contribute to the structural maintenance of LCCs and other potential roles in chromatin regulation, besides their roles as epigenetic marks for transposons, await further investigation in future studies.

## Methods

### Plasmids and antibodies

For the expression of dinoflagellate TET/JBP homologues in *Escherichia coli* BL21 (DE3), codon optimized AmpTJ1 and CryTJ1 genes were individually synthesized and cloned into the pET28a plasmid by Tsingke Biological Technology Co., Ltd. (Beijing, China), yielding pET28a-AmpTJ1 and pET28a-CryTJ1 plasmids, respectively. The CryTJ1 mutant (HRD to ARA mutation, CryTJ1m) was generated by site-directed mutagenesis to form pET28a-CryTJ1m plasmid. To construct a stable HEK293T cell line expressing CryTJ1, CryTJ1 with three N-terminal SV40 nuclear localization signals (3×NLS) and a C-terminal FLAG-tag was cloned into the lentiviral pLVX-Tetone-Puro vector (Tetone), producing the plasmid pLVX-3×NLS-CryTJ1-Flag-Tetone-Puro. To heterogeneously express CryTJ1 and CryTJ1m in budding yeast, we fused the FLAG-tagged 3×NLS-CryTJ1 and 3×NLS-CryTJm with GFP (green fluorescent protein) gene individually and cloned into a yeast expression plasmid pESC-Leu, resulting in pESC-3×NLS-CryTJ1-GFP-Flag and pESC-3×NLS-CryTJ1m-GFP-Flag vectors. The antibodies used were anti-5hmU antibody (ab19735, Abcam), Anti-Flag antibody (F1804, Sigma), anti-DYKDDDDK antibody (66008-3-Ig, Proteintech), anti-GAPDH antibody (60004-1-Ig, Proteintech) and goat anti-mouse IgG secondary antibody (L3032, Signalway Antibody).

### Cell culture, cell cycle synchronization and flow cytometry analysis

The cultures of dinoflagellates *Amphidinium carterae* and *Symbiodinium* sp. (from Shanghai Guangyu Biotechnology Co., Ltd., China) were maintained in seawater-based f/2 medium at 23°C, under illumination at 50 µmol photons m^−2^ s^−1^ on a 12 hour : 12 hour light-dark cycle. The cell counts of *A. carterae* were measured using a Countess TM II FL cell counter (Thermo Fisher). Dinoflagellate *Crypthecodinium cohnii* ATCC 30556 (from the American Type Culture Collection) was cultured in A2E6 medium without light at 23°C. HEK293T cells were normally grown in DMEM (Gibco) supplied with 10% fetal bovine serum (FBS, BI) and 1% penicillin-streptomycin (Thermo Fisher) at 37°C with 5% CO_2_ in a humidified incubator.

For synchronization, exponentially grown *A. carterae* cells were firstly diluted to 5 × 10^4^ cells/ml, followed by 2 days’ darkness, and then returned to the normal 12 hour : 12-hour light-dark cycle. After one day incubation and at the beginning of another light-dark cycle, this synchronized culture was collected in different time points at T0, T4, T8, T10, T11, T12, T14, T16, T20 and T24 during the 24-hour period.

*A. carterae* cell samples for cell cycle examination were harvested by centrifugation at 3000 rpm for 10 minutes. Cells were fixed with pre-cooled 70% ethanol on ice bucket for at least one hour. After fixation, cells were resuspended with cold methanol and stored for 24 hours to facilitate pigment removal. The cell pellets were then washed and resuspended in 980 µl of 1 ×Phosphate-buffered saline (1 ×PBS, pH 7.4), followed by addition of 20 µl of 0.5 mg/ml RNase A (NEB) and incubation at 37°C for 1 hour. Finally, each sample was stained with 25 µg/ml propidium iodide and incubated at room temperature for 30 minutes in darkness. Samples were then transferred to 5ml BD Falcon tubes and examined with a BD FACSCanto SORP Flow Cytometry System (BD Biosciences). The 70% ethanol fixed Human iPSC sample was a kind gift from Prof. Tian Ruilin, SUSTech.

### Treatment of *A. carterae* cells with [^15^N_2_] dT or 2,4-Pyridinedicarboxylic acid hydrate

To determine the potential source of 5hmU in *A. carterae*, exponentially growing cells were diluted at a density of 1 ×10^5^ cells/ml and incubated in darkness for two days. Subsequently, 50 µM of [^15^N_2_]-labeled dT (Silantes) were adding to the cell culture. At daily intervals thereafter, cells were harvested via centrifugation, followed by gDNA extraction using the CTAB method described later.

To inhibit the synthesis of 5hmU in *A. carterae*, cells in the exponential stage (two biological replicates) were diluted to a concentration of 1 × 10^5^ cells/ml in a 6-well plate and treated with 0.5 mM 2,4-pyridinedicarboxylic acid hydrate (2-4P) in f/2 medium. After 3-days’ incubation, cells were collected to perform total RNA and gDNA extraction. The total RNA and gDNA were then subjected to RNA-sequencing and LC-MS/MS analysis respectively.

### Genomic DNA extraction

Exponential-phase *A. carterae* cells (∼2 ×10^7^ cells) were collected at 3000 rpm for 10 minutes and the genomic DNA (gDNA) were subsequently extracted via a CTAB method. Briefly, cell pellets were resuspended in 600 µl of preheated CTAB extraction buffer (100 mM Tris pH 8.0, 20 mM EDTA pH 8.0, 2 M NaCl, 2% (w/v) CTAB, 2% (w/v) Polyvinylpyrrolidone. Next, 10 µl of 20 mg/ml RNase A (NEB) was added, and the mixture was incubated at 37°C for 2 hours, followed by addition of 20 µl of Proteinase K (NEB) and another incubation at 65°C overnight. Equal volumes of phenol : chloroform : isoamyl alcohol (25 : 24 : 1) (Solarbio Life Sciences, Beijing, China) were added. After centrifugation, the upper phase was transferred to a new Eppendorf tube, mixed with 2/3 volume of isopropanol and then centrifuged at 13000 rpm for 10 minutes. The pellet was then washed with 700 µl of 75% ethanol and 80% ethanol sequentially. Finally, the DNA was eluted in 20 µl of ddH_2_O. For gDNA extraction from *Symbiodinium* sp. and *C. cohnii*, exponential-phase cells were resuspended in 600 µl of CTAB extraction buffer and lysed with acid-washed glass beads (Sigma) using a FastPrep-24 5G (MPBio) homogenizer for two cycles (6m/s, 40s on and 60s off). The remaining steps were identical with the gDNA extraction of *A. carterae* cells. For HEK293T cells, gDNA extraction was performed using the FastPure Cell/Tissue DNA Isolation Mini Kit (Vazyme Biotech, Nanjing, China).

### Sample preparation for LC-MS/MS

For LC-MS/MS analysis, 200ng of gDNA was firstly denatured at 100°C for 10 minutes and cooled immediately on ice for 5 min. Then gDNA was digested into nucleosides using the nuclease P1 (NEB) and Antarctic Phosphatase (NEB) following the manufacturer’s instructions. The enzymatic reaction was stopped with 75µl of methanol. Following centrifugation, the supernatant was freeze-dried and resuspended in 50 µl of ddH_2_O.

LC-MS/MS analysis was performed on a Q Exactive high-resolution benchtop quadrupole Orbitrap mass spectrometer under positive-ion mode using an Agilent XDB-C18 column. The samples were run in a mobile phase composed of buffer A (water with 0.1% formic acid) and 2% to 98% gradient of buffer B (methanol in 0.1% formic acid). Nucleosides were identified by their retention time and specific ion mass transitions: 5hmU, 259.0924 to 125.0349; dT, 243.0975 to 127.0505; 5mC, 242.1135 to 126.0665; 5hmC, 258.1084 to 142.0613; 6mA, 266.1247 to 150.0800; 5fU, 279.0587 to 163.0113; [^15^N_2_]5hmU, 261.0865 to 127.0290; and [^15^N_2_] dT, 245.0916 to 129.0446.

### RNA sequencing

Total RNA from *A. carterae* was extracted using RNAzol®RT (MRC) following the manufacturer’s instruction. For genome annotation, the short-read RNA-seq data and long-read transcriptome of *A. carterae* were sequenced by the Beijing Genomics Institute (BGI, Shenzhen, China) on the DNBSEQ PE150 and PacBio Sequel II platforms respectively, according to manufacturer’s instructions. For 2-4P treatment of *A. carterae* cells, the extracted total RNA was first treated with Ribo-off® rRNA depletion kit (N409, Vazyme Biotech, Nanjing, China) to remove rRNA using self-designed antisense oligos according to a previous study ^40^. Then the rRNA-depleted RNA samples were sent to either Beijing Genomics Institute (BGI, Shenzhen, China) or Novogene Company (Beijing, China) for stranded RNA-seq libraries construction and sequencing.

### Genome survey

gDNA extracted from *A. carterae* cells was sent to Beijing Genomics Institute (BGI, Shenzhen, China) and sequenced by the DNBSEQ PE150 platform. The raw sequencing data was filtered with SOAPnuke (V1.6.5) to remove low-quality reads, adapter contaminants, and PCR duplicates. The resulting high-quality reads were then used to count the 21-kmer frequency. Subsequently, GenomeScope was used to model the 21-kmer spectrum and estimate genome size.

### Genome sequencing and *De novo* genome assembly

Exponentially growing *A. carterae* cells were sent to Beijing Genomics Institute (BGI, Shenzhen, China) for gDNA extraction and Hi-C library construction. All Pacbio HiFi libraries were sequenced on the Pacbio Sequel II platform (BGI, Shenzhen, China) with CCS mode, achieving an approximate depth of 30 × based on the estimated genome size. Standard Hi-C library was sequenced on the DNBSEQ platform (BGI, Shenzhen, China).

The genome of *A. carterae* was firstly assembled using Pacbio HiFi sequencing reads and further refined with Hi-C technology to determine contig position on scaffolds. A contig-level assembly of the resulting PacBio CCS reads with default parameter was obtained using Hifiasm software. Assembled contigs were further joined into scaffolds at pseudo-chromosome level using Hi-C sequencing data with Juicer (default parameters) and 3D-DNA (-m haploid -q 0 -r 0).

### Genome annotation

Repeat sequences were annotated using a combination of homology-based and *de novo* methods. RepeatMasker (v4.0.7) and RepeatProteinMask (v4.0.7) were used to identify known repeat sequences through searching against RepBase v21.12 library. On the other hand, in the *de novo* step, *de novo* repeat sequence libraries were established using RepeatModeler and LTR_FINDER v1.06, followed by *de novo* predication with Repeatmasker program. Additionally, Tandem Repeats Finder v4.09 was used to detect tandem repeats in the genome.

Protein-coding genes were identified using a combination of homology-based, *de novo* and transcriptome-based strategies. For homology-based approach, genes of 7 dinoflagellate species including *Amphidinium carterae* CCMP1314 (HBNO00000000.1)*, Amphidinium gibbosum* ^41^, *Fugacium kawagutii* ^24,42^, *Polarella glacialis_*CCMP1383 ^24^, *Polarella glacialis* CCMP2088 ^24^, *Symbiodinium microadriaticum* ^19^, and *Symbiodinium minutum* ^43^were downloaded from public database, and search against our assembled *A. carterae* genome to obtain gene models. For RNA-seq data alignment method, short reads from DNBSEQ were mapped to the genome assembly of *A. carterae* for identifying splice sites with HISAT2 and then assembled via Stringtie. Meanwhile, the long-read transcriptome was used to search against the *A. carterae* genome using Exonerate. For *de novo* predication, 2000 genes with Annotation Edit Distance values < 0.05 and complete structures from the above predicted genes were randomly selected to train models for the tools AUGUSTUS and SNAP. Then, we combined the gene predication results from *de novo* prediction, homology-based and transcriptome-based annotations and used MAKER-P software to obtain the integrated consensus gene models. Finally, gene function annotation was obtained with BLAST search against reference databases including SwissProt, TrEMBL, KEGG, InterPro, NR, KOG, and GO. Noted that for genome characteristics comparison, we also downloaded assembled genomes from dinoflagellates including *Symbiodium natans* CCMP2548 ^22^, *S. tridacnidorum* CCMP2592 ^22^, *Cladocopium goreaui* SCF055 ^44^ and *Prorocentrum cordatum* CCMP1329 ^25^.

### Phylogenetic analysis of Tet/JBP family proteins

Tet/JBP protein homologues in different dinoflagellates were retrieved from *C. cohnii* ATCC 30556 transcriptome (GFIV00000000.1), our *A. carterae* genome and the marine microbial eukaryote transcriptome sequencing project (MMETSP) using the TET_JBP domain (PF12851) as a query for iterative HMMER searches. Phylogenetic analysis was conducted using built-in IQ-TREE program in TBtools with 1000 ultrafast bootstraps ^45^. The resultant phylogenetic tree was visualized with the Evolview platform ^46^. Multiple sequence alignments of the Tet/JBP homologues were performed with NCBI online tool COBALT ^47^ and visualized via the ESPript web server ^48^.

### Characterization of TJ1 enzyme activities *in vivo*

Plasmids pET28a, pET28a-AmpTJ1, pET28a-CryTJ1 and pET28a-CryTJ1m were firstly transformed into *E. coli* BL21 (DE3) cells individually. These transformants were normally grown at 37 °C in a Luria-Bertani (LB) medium supplemented with 50 µg/mL of kanamycin. When *E. coli* BL21 (DE3) cells carrying each plasmid grown at an optical density of 0.6-0.8, isopropyl β-D-1-thiogalactopyranoside (IPTG) was added at a final concentration of 0.5 mM and incubated with the cell culture at 16 °C for 16 hours to induce protein expression. After centrifugation, a portion of the cell pellets were mixed with 6 × protein loading buffer, analyzed by SDS-PAGE, and visualized using Coomassie blue staining. On the other hand, genomic DNA was isolated using the TIANamp Bacteria DNA Kit (TIANGEN Biotech Co., Ltd., China). LC-MS/MS technique was employed to detect DNA base modifications on the extracted *E. coli* gDNA.

### Expression and purification of recombinant CryTJ1 protein

After IPTG-induction of protein expression for *E. coli* BL21 (DE3) strain containing pET28a-CryTJ1 plasmid, the cells were centrifuged at 6000 rpm for 20min at 4°C and washed with pre-cooled 1 ×PBS. Cell pellets were resuspended in lysis buffer (20 mM Tris-HCl (pH 8.0), 300 mM NaCl, 15 mM imidazole, 10% glycerol and 0.25% Triton X-100) and lysed using a high-pressure homogenizer (AH-NANO, ATS, China) with pressure about 800 ∼ 900 bar. The lysates were then centrifuged at 20000 rpm for 40 min and the resulting supernatant was incubated with Ni-NTA agarose beads (Changzhou Smart-Lifesciences Biotechnology Co., Ltd., China) at 4°C. After washing with wash buffer (20 mM Tris-HCl (pH 8.0), 300 mM NaCl, 50 mM imidazole, 10% glycerol), rCryTJ1 protein was eluted with elution buffer (20 mM Tris-HCl (pH 8.0), 150 mM NaCl, 300mM imidazole) and examined by Coomassie blue staining. The final rCryTJ1 was stored in a buffer containing 50 mM HEPES-NaOH (pH 8.0), 100 mM NaCl, and 10% glycerol.

### *In vitro* enzyme activity assay of rCryTJ1

Purified rCryTJ1 (500ng) was combined with 250ng of HEK293T gDNA in a reaction buffer consisting of 50 mM HEPES-NaOH (pH 8.0), 100 mM NaCl, 75 µM Fe(NH_4_)_2_(SO_4_)_2_, 2 mM ascorbate, 1 mM α-ketoglutarate and 1 mM ATP, followed by incubation at 28°C overnight. For the control experiment, an equivalent quantity of rCryTJ1 was inactivated at 100°C for 10 minutes before being mixed with 250 ng HEK293T gDNA, and then used to carry out reaction under the same condition.

To assess the impact of different metal ions on rCryTJ1 activity, we omitted the metal ions in the same reactions mentioned above, which were then individually supplemented with 75 µM of Fe(NH_4_)_2_(SO_4_)_2_, MgCl_2_ and MnCl_2_. Experiments were also conducted in the absence of ATP and ascorbate. Meanwhile, the activity of rCryTJ1 toward ssDNA from HEK293T was tested.

To understand the effect of nucleosome wrapping on the catalytic activity of rCryTJ1, recombinant mononucleosome (81070, Active motif) comprising 167 bp of Widom 601 DNA and the corresponding naked dsDNA were used as substrates respectively. They were individually combined with rCryTJ1 in the aforementioned reaction buffer and then incubated at 28°C overnight. After the enzymatic reaction, each sample was treated with nuclease P1 (NEB) and Antarctic Phosphatase (NEB) to digest the gDNA and then processed for LC-MS/MS analysis.

### Oxidation of 5hmU to 5fU with ACT^+^BF_4^−^_

A recent study reported that ACT^+^BF_4^−^_, a stable, water soluble and environmentally friendly chemical, was able to fully oxidize 5mC into 5hmC at dsDNA level under a much milder condition when compared with ruthenium-based oxidation approaches ^29^. Thus, we first tested whether ACT^+^BF_4^−^_ could also be utilized to completely oxidize dsDNA 5hmU into 5fU. In brief, gDNA extracted from *A. carterae* cells was fragmented using the Bioruptor Pico sonication device (Diagenode) to reach a size range of 200-400 bp. Following fragmentation, the DNA was purified using the DNA Clean & Concentrator-5 Kit (Zymo Research) according to the manufacturer’s protocol. An aliquot of this purified DNA (∼500 ng) was then incubated in a 25µl solution comprising 50 mM phosphate buffer (pH7.5) and 50 mM ACT^+^BF4^−^ at 37°C for either 4 or 8 hours. After oxidation, the DNA was further purified using the Oligo Clean & Concentrator Kit (Zymo Research). The purified samples were then ready for subsequent treatment.

### 5hmU-DIP and 5fU-DIP library construction and sequencing

*A. carterae* gDNA was first sonicated to 200-400 bp using the Bioruptor (Diagenode). A portion of sonicated DNA were retained as input sample. Both input and the remaining sonicated *A. carterae* gDNA underwent end-repair using the VAHTS® Universal DNA Library Prep Kit for Illumina V3 (ND607, Vazyme Biotech, Nanjing, China). Subsequently, adapters from the VAHTS DNA Adapters Set 2 for Illumina (N802, Vazyme Biotech, Nanjing, China) were ligated as the manufacturer’s instructions. The product was purified using 0.8× VAHTS DNA Clean Beads and washed twice with an 80% fresh-prepared acetonitrile. For the 5hmU-DIP, the fragmented gDNA (∼120ng) was directly incubated with (∼1µl) anti-5hmU antibody (Abcam, ab19735). Then antibody and DNA mixture were incubated in 1 ×IP buffer (10 mM Na-Phosphate pH 7.0, 140 mM NaCl, 0.05 % Triton X-100) overnight at 4°C with gentle rotation. After incubation, 20ul of prewashed Protein A/G beads (Thermo Fisher Scientific) were added to each tube for 2 hours of incubation at 4°C. After binding the DNA-antibody complexes to the beads, they were washed five times with 1× IP buffer. The beads were then resuspended in 200µl of protease K digestion buffer (50 mM Tris, pH 8.0, 10 mM EDTA, 0.5 % SDS), treated with 2 µl of protease K, and incubated at 50°C for 2 hours. After digestion, the supernatant was purified using the Oligo Clean & Concentrator Kits (Zymo Research). The resultant DNA was eluted in 15 µl ddH_2_O, and its concentration was determined using the Equalbit 1 × dsDNA HS Assay Kit (Vazyme Biotech, Nanjing, China). DNA libraries were then amplified from enriched DNA and the input samples using the VAHTS® Universal DNA Library Prep Kit for Illumina V3 and checked using a BiOptic Qsep100 Bio-Fragment Analyzer with the standard S2 cartridge. For the parallel 5fU-DIP, ACT^+^BF_4^−^_ oxidation was performed firstly to fully convert gDNA 5hmU into 5fU, and equal amounts of oxidized gDNA were then treated with the same quantity of anti-5hmU antibody. The other steps were identical to 5hmU-DIP procedures. Finally, all constructed libraries underwent Illumina short-read sequencing on the Hiseq PE150 platform by Novogene Biotech Co., Ltd (China). On the other hand, we also used the same procedures to construct a 5hmU-DIP library based on the isolated gDNA of *Symbiodinium* sp.. Note that its 5hmU-DIP library was sequenced on the MGISEQ-2000 platform (BGI, Shenzhen, China).

### 5hmU-chemical-seq

For 5hmU-chemical-seq, we utilized both commercial adapters (N802, VAHTS DNA Adapters Set 2 for Illumina, Vazyme Biotech, Nanjing, China) and modified adapters inspired by a previous study ^13^. The modified adapter includes a 5’-end modified oligodeoxynucleotide (ODN 1) bearing an amino group (NH2) attached via a six-carbon (C6) spacer (NH_2_-C6), with a sequence of 5’-NH_2_-C6-GAATGATACGGCGACCACCGAGATCTACACTCTTTCCCTACACGACGCT CTTCCGATCT-3’. Additionally, we employed a 3’-phosphorylated ODN with a sequence 5’-phosphate-GATCGGAAGAGCACACGTCTGAACTCCAGTCACNNNNNNATCTCGTATGCCGTCTT CTGCTTG-phosphate-3’. In the present study, NNNNNN was corresponding to ATCACG (ODN 2) and GATCAG (ODN 3) respectively. ODN 1 was individually annealed with ODN 2 and ODN 3 in a solution of 50 mM NaCl and 10 mM Tris-HCl (pH 7.0), resulting in the final adapters 5N3A3 and 5N3A4, respectively.

The remaining steps were basically similar to previous procedures ^13^. In brief, *A. carterae* gDNA samples (∼2000 ng) were fragmented and subjected to end-repair. For input sample, a portion of DNA was ligated with adapters from the VAHTS DNA Adapters Set 2 for Illumina (N802, Vazyme Biotech, Nanjing, China). On the other hand, the recovered end-repaired gDNA was ligated to the pre-annealed adapters 5N3A3 and 5N3A4 using the VAHTS® Universal DNA Library Prep Kit for Illumina V3 (ND607, Vazyme Biotech, Nanjing, China). Following ligation, the DNA samples were purified with 0.8× VAHTS DNA Clean Beads (N411, Vazyme Biotech, Nanjing, China) and washed twice with freshly prepared 80% acetonitrile. The samples were then oxidized using ACT^+^BF4^−^.

The resulting oxidized DNA samples (∼300ng) were mixed with 20 mM phosphate buffer (pH 6) and 5.1 mM (+)-biotinamidohexanoic acid hydrazide (Sigma-Aldrich), and incubated at 40°C for 4 hours. After incubation, these mixtures were purified using the Oligo Clean & Concentrator Kits (Zymo Research). Biotinylated DNA samples were added to streptavidin magnetic beads (Thermo Fisher) that had been pre-washed using a binding buffer composed of 10 mM Tris (pH 7.5), 1 mM EDTA, 2M NaCl, and 0.1% (v/v) Tween 20. This mixture was rotated gently at 4°C for 30 minutes. Then the samples were washed five times with the binding buffer on a magnetic rack. DNA was eluted with elution buffer (0.05% (v/v) hydroxylamine, 5 mM P-Anisidine, 0.1% tween 20 in 50 mM phosphate buffer, pH 6) at 40°C with vigorous shaking at 1400rpm for 30 minutes using a thermal mixer (Thermo Scientific). The eluted DNA and previous input samples were then amplified using the PCR Primer Mix 3 for Illumina and the VAHTS HiFi Amplification Mix 2 (ND607, Vazyme Biotech, Nanjing, China). These libraries were then sequenced on the Hiseq-PE150 platform by Novogene Biotech Co., Ltd.

### Stable transfection of HEK293T cells

The cell transfection was carried out according to the previous procedures ^49^. Briefly, when HEK293T cells in a 6-well plate reached approximately 70% confluence, they were co-transfected with lentivirus packaging plasmids including psPAX2 and pMD2.G, and target plasmid pLVX-3×NLS-CryTJ1-Flag-Tetone-Puro or empty vector pLVX-Tetone-Puro using Lipofectamine 3000 Transfection Reagent (Invitrogen). At 48 hours post-transfection, lentiviral supernatant was harvested, aliquoted and stored at −80°C. Meanwhile, the remaining cells were sub-cultured onto a new 6-well plate and screened with 2µg/ml puromycin. After a week of selection with puromycin, cells were treated with three different concentrations including 0 ng/ml, 100ng/ml and 1000ng/ml of Doxycycline (Dox) to induce CryTJ1 expression. Three days after doxycycline treatment, the cells were collected and subjected to immunoblotting, LC-MS/MS and immunofluorescence analysis.

### Immunoblotting

Cell lysates from HEK293T cells were directly lysed with 2 ×SDS-PAGE (sodium dodecyl sulphate polyacrylamide gel electrophoresis) protein loading buffer. The total protein extraction from budding yeast was conducted according to a previous standard protocol ^50^. The lysate was boiled at 100°C for 5 minutes and then subjected to separation with SDS-PAGE, which were subsequently transferred onto PVDF (polyvinylidene difluoride; Merck Millipore) membrane. After transfer, the membranes were blocked with 5% milk in TBST (20 mM Tris (pH 7.4), 150 mM NaCl, and 0.2% Tween-20) for 1 hour. The membranes were then incubated overnight at 4°C with anti-DYKDDDDK antibody (66008-3-Ig, 1:2000 dilution, Proteintech) in Western Primary Antibody Dilution Solution (Beyotime Biotech, Shanghai, China). After washing the membranes with TBST, they were incubated with a goat anti-mouse IgG secondary antibody (L3032, Signalway Antibody LLC, Maryland, USA) for 1 hour at room temperature and washed again with TBST. The membranes were then developed using the Clarity Western ECL Substrate (BIO-RAD). For the loading control, the same membrane was subsequently probed with anti-GAPDH antibody (1:2000 dilution, 60004-1-Ig, Proteintech) overnight at 4°C.

### Immunofluorescence microscopy

Immunofluorescence was carried out using a previously established protocol ^51^. In brief, HEK293T cells expressing CryTJ1 were seeded on glass cover-slip in 24-well plates. After Dox induction, the cells on cover-slip were fixed with 4% paraformaldehyde for 15 minutes and then rinsed three times with 1 × PBS. Permeabilization was conducted by treating the cells with 0.5% Triton X-100 in 1 × PBS for 20 minutes, followed by three washes with 1 × PBS for 3 minutes. To prevent non-specific antibody binding, cells were blocked with a mixture of 2% bovine serum albumin (BSA, Beyotime Biotech, Shanghai, China) and 2% normal donkey serum in 1 × PBS for 30 minutes. The cells were then incubated with a 1:500 dilution of anti-Flag antibody (F1804, Sigma-Aldrich) for 2 hours at room temperature. After wash, the cells were treated with Alexa Fluor 488 Goat Anti-Mouse IgG antibody (L3032, 1:1000 dilution, Thermo Fisher) for 30 minutes at room temperature. Following another wash with PBST (1 ×PBS and 0.2 % Tween-20), cell nuclei were stained using DAPI Fluoromount-G (SouthernBiotech). Images were finally captured using a Zeiss LSM980 confocal microscope.

### Yeast transformation

The transformation was performed in a leucine auxotroph budding yeast strain (BY4741, a generous gift from Professor Hai Rao, SUSTech) using a lithium acetate-based protocol as previously described ^52^. After transformation with the pESC-3×NLS-CryTJ1-GFP-Flag and pESC-3×NLS-CryTJ1m-GFP-Flag plasmids individually, yeast cells were cultured in a raffinose containing basic medium overnight. On the next day, a 1:5 dilution of this overnight culture was grown in 8 mL same medium at 30°C with rotation until exponential growth was attained. Then galactose was added at a final concentration of 2% (wt/vol) and incubated with the cell culture for another 24 hours to induce protein expression. The expression of the target proteins was verified by western blotting and confocal microscopic observation of the green fluorescent protein (GFP) signal. Hoechst 33342 at a final concentration of 5µg/ml was used to stain the yeast nuclei. The yeast gDNA was extracted by YeaStar Genomic DNA Kit (Zymo Research) and then proceeded to LC-MS/MS analysis.

### Whole genome bisulphite sequencing (WGBS)

*A. carterae* gDNA was sent to Novogene Biotech Co., Ltd (China) for WGBS libraries construction according to a standard protocol. Briefly, gDNA was spiked with unmethylated lambda DNA and sheared to average 200 bp size, followed by bisulphite treatment using the Accel-NGS® Methyl-Seq DNA Library Kit (Swift Biosciences). Following this treatment, the DNA was subjected to end-repair, adapter ligation, and PCR amplification. Finally, libraries were sequenced on a Hiseq-PE150 platform (Novogene Biotech Co., Ltd, China).

For WGBS analysis, bisulfite-treated reads were filtered to remove adaptors and low-quality bases. The resulting clean reads were aligned to the *A. carterae* genome (Acart.genome.v1) using BS-Seeker2 (v2.1.8, --aligner=bowtie2 --local). The CGmap output file generated by BS-Seeker2 for methylation calling from the mapping result was utilized in downstream analysis.

### Bioinformatic analyses

Raw DIP-seq and 5hmU-chemical-seq data were first processed using TrimGalore (version 0.6.7) to remove adapters and low-quality bases (-q 25). The cleaned sequence reads were then mapped to the *de novo* reference genome (Acart.genome.v1) using Bowtie2 local alignments. Resultant SAM formatted files were converted to the BAM format using Samtools. Enriched regions were identified with MACS2 (-g 1.2e9 -q 0.05). Motif discovery was performed using MEME (-mod zoops -minw 4 -maxw 10 -nmotifs 10 -dna) and the genomic distribution of peaks was analyzed using Bedtools.

## Acknowledgments

We would like to thank Prof. Chaoxing Liu at Sun Yat-sen University and Prof. Hongjie Shen at Fudan University for their advice in experimental design. The authors would also like to acknowledge the technical support from Hua Li and Lin Lin at SUSTech CRFT. This work was supported by Center for Computational Science and Engineering at Southern University of Science and Technology.

This work was supported by National Key Research and Development Program of China (2022YFC2702705), National Natural Science Foundation of China (32170604 to H.C.). This work was also supported by Pearl River Recruitment Program of Talents (2021QN02Y122) and Department of Health of Guangdong Province (B2021032) to H.C., Shenzhen Key Laboratory of Gene Regulation and Systems Biology (Grant No. ZDSYS20200811144002008) from Shenzhen Innovation Committee of Science and Technology and Funding for Scientific Research and Innovation Team of The First Affiliated Hospital of Zhengzhou University (ZYCXTD2023004).

## Author contributions

C.P.L. and H.C. designed and conceived the experiments. C.P.L., Y.L. (Ying LI) and H.Z. performed most of the experiments. Under the supervision of J.W.X. and W.Q.X., Y.C.W., Y.L. (Ying LI) and R.X.M analyzed the NGS data. X.Y.S., Y.Y.Z. and N.L. assisted in HPLC-MS/MS experiment. H.D.H and Y.L. (Yue LI) contributed to the enzymatic analysis of TJ1. C.P.L. and H.C. wrote the manuscript with the inputs from other authors. All authors have read and approved the final manuscript.

## Supplementary Figure Legends

**Figure S1.**
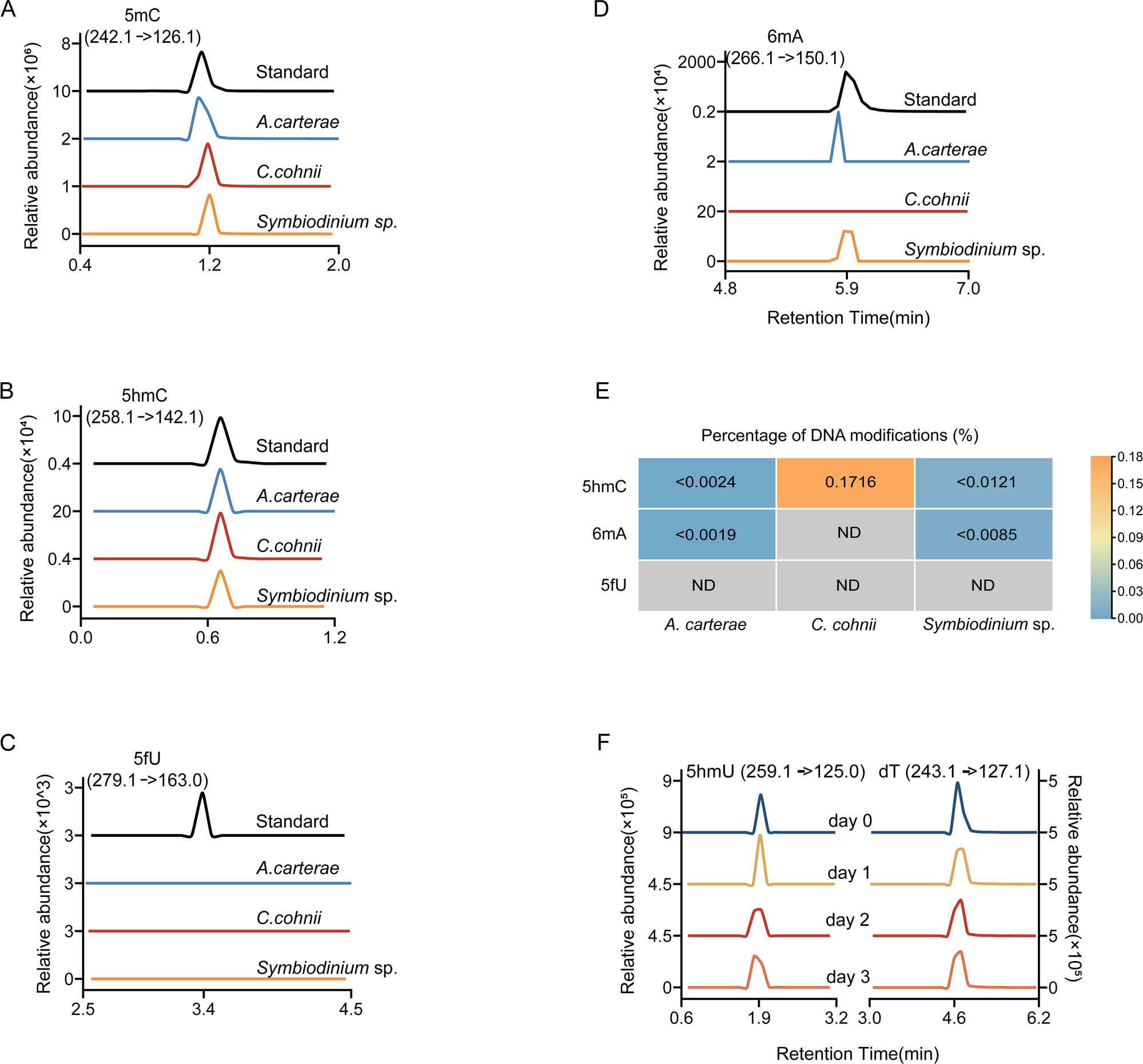
Analysis of common DNA modifications in *A. carterae* genomic DNA. (A-D) LC-MS/MS analysis of DNA modifications including 5mC (A), 5hmC (B), 5fU (C), and 6mA (D) within different dinoflagellates including *A. carterae*, *C. cohnii*, and *Symbiodinium* sp.. (E) LC-MS/MS quantification of 5hmC, 6mA, and 5fU in the gDNA of the three dinoflagellate species. Three biological replicates were used. ND: Not Detected. (F) DNA spectra of 5hmU and dT in extracted gDNA of *A. carterae* cells incubated with nitrogen stable isotope-labeled thymidine ([^15^N_2_]dT).

**Figure S2.**
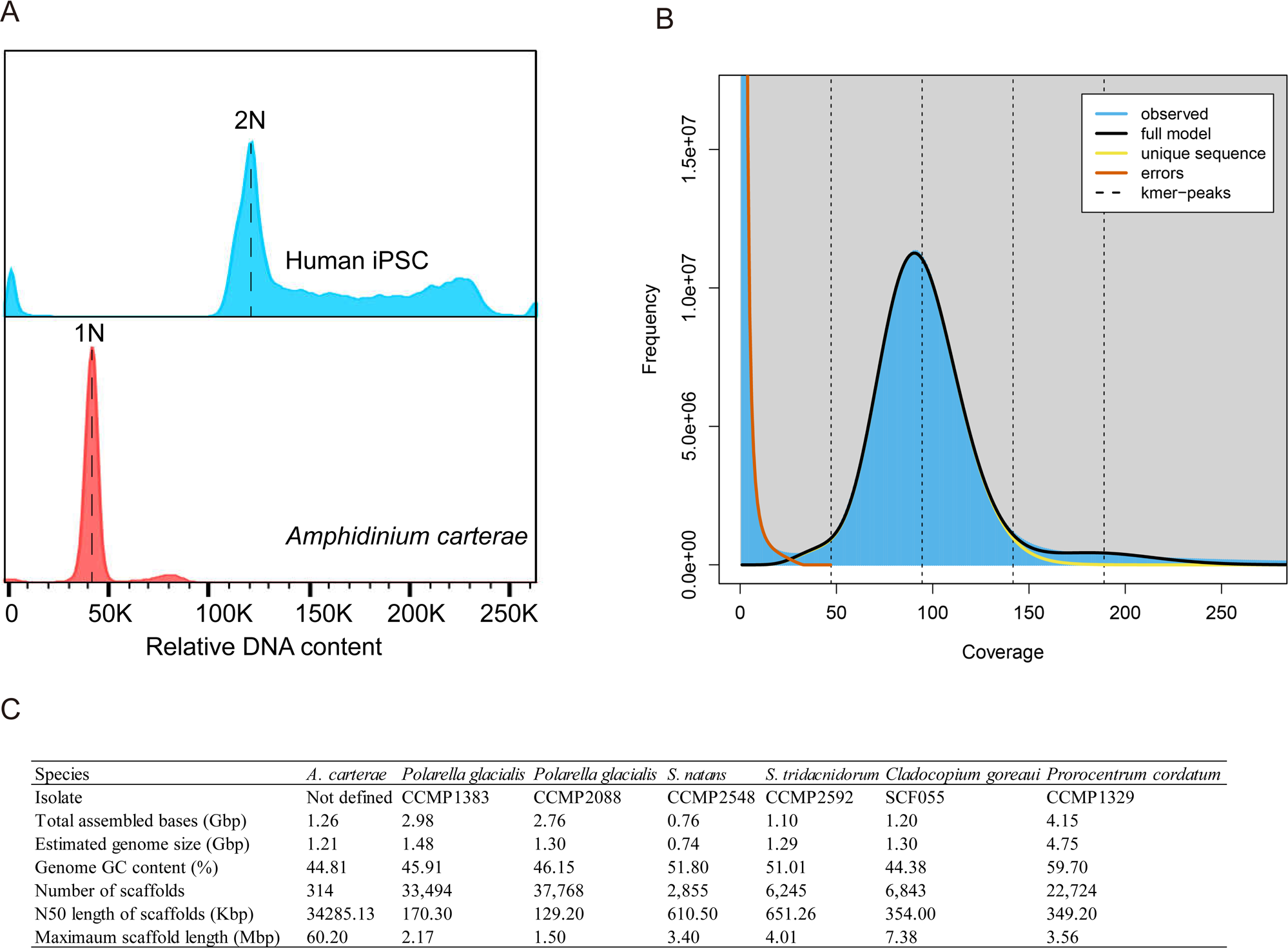
Genome size estimation and the features of *A. carterae* genome. (A) Flow cytometry analysis of the DNA content of human induced pluripotent stem cells (IPSC) and dinoflagellate *A. carterae.* (B) GenomeScope profile for *A. carterae* based on 21-mers using illumina paired-end reads. (C) A comparison of genome assembly characteristics between *A. carterae* and other dinoflagellates including *Polarella glacialis* CCMP1383 and CCMP2088, *Symbiodium natans* CCMP2548, *S. tridacnidorum* CCMP2592, *Cladocopium goreaui* SCF055 and *Prorocentrum cordatum* CCMP1329.

**Figure S3.**
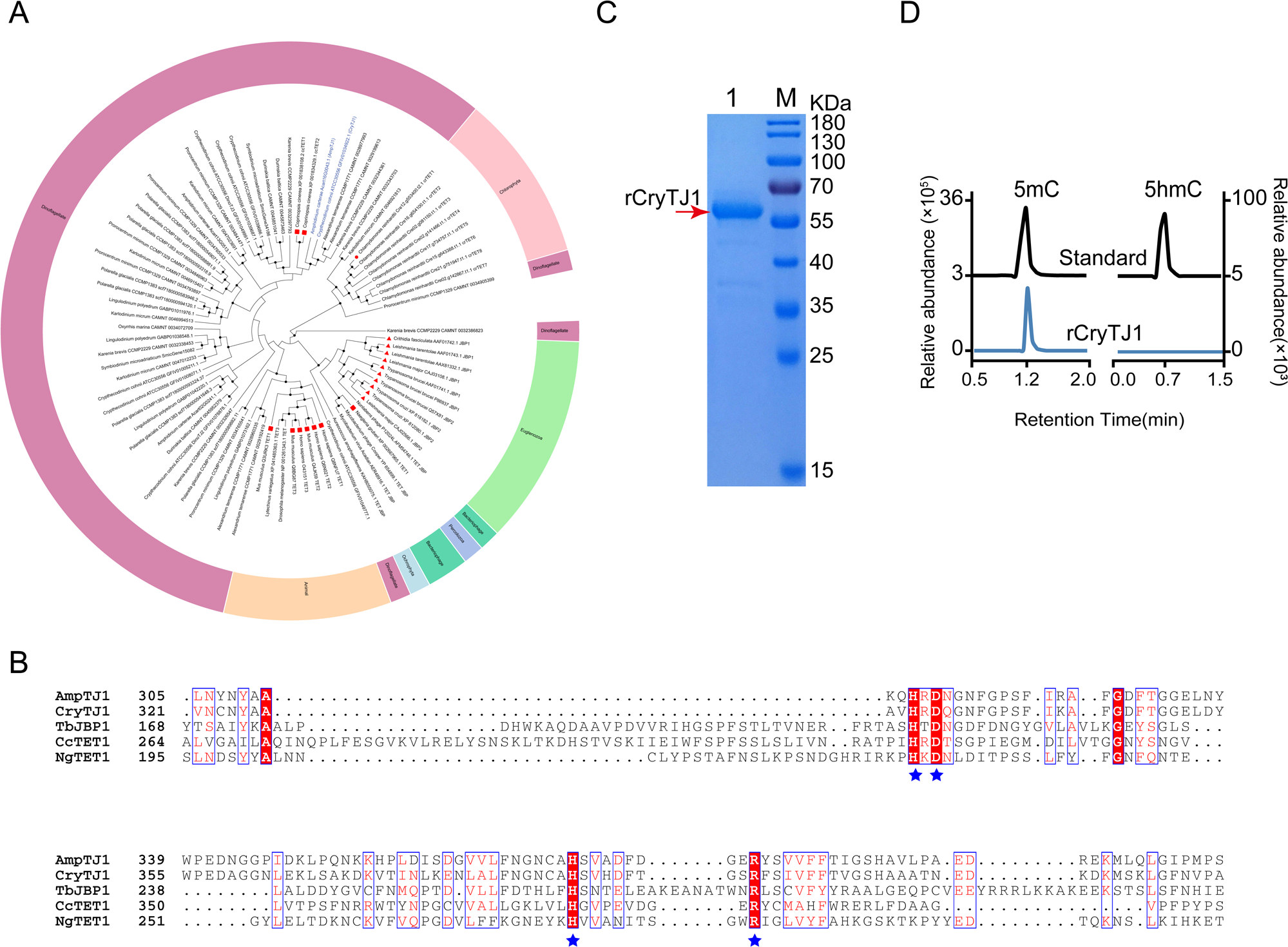
Identification and characterization of putative dinoflagellate Tet/JBP family protein homologues. (A) Phylogenetic analysis of Tet/JBP proteins from different dinoflagellates and other representative eukaryotes. Rectangle, Triangle, and Circle represent known TET/JBP homologues with enzyme activities to catalyze conversion of 5mC to 5hmC, thymidine to 5hmU, and 5mC to 5gmC (5-glyceryl-methylcytosines) respectively. (B) Multiple sequence alignment of CryTJ1, AmpTJ1, and other representative TET/JBP proteins. Star, highlights the HXD and H•••R motifs that are critical for binding cofactors Fe^2+^ and 2-oxoglutarate (2OG). (C) SDS-PAGE showing the expected size (∼59.5 kDa) of the purified rCryTJ1 proteins. (D) Purified rCryTJ1 couldn’t convert 5mC of HEK293T gDNA into 5hmC at the dsDNA level.

**Figure S4.**
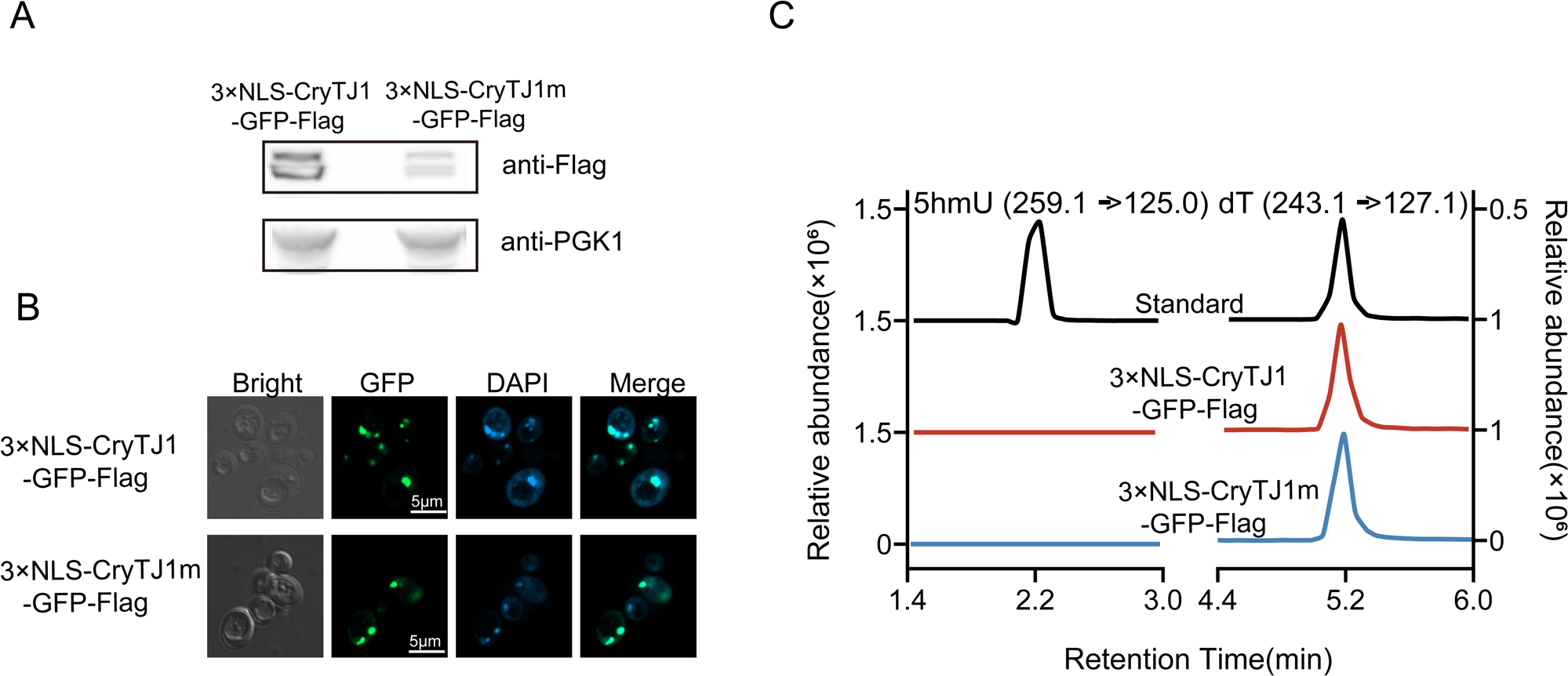
Heterogeneous expression of CryTJ1 in budding yeast cells cannot lead to 5hmU production. (A) Immunoblots of budding yeast cells expressing either pESC-3×NLS-CryTJ1-GFP-Flag (3×NLS-CryTJ1-GFP-Flag) or pESC-3×NLS-CryTJ1m-GFP-Flag (3×NLS-CryTJ1m-GFP-Flag) after galactose induction. PGK1 was used as loading controls. (B) Confocal micrographs indicating colocalization between the GFP signals fused with CryTJ1 or CryTJ1m (green) and Hoechst staining (blue). (C) LC-MS/MS analyses results showing no generation of 5hmU after galactose induction for different pESC-Leu constructs.

**Figure S5.**
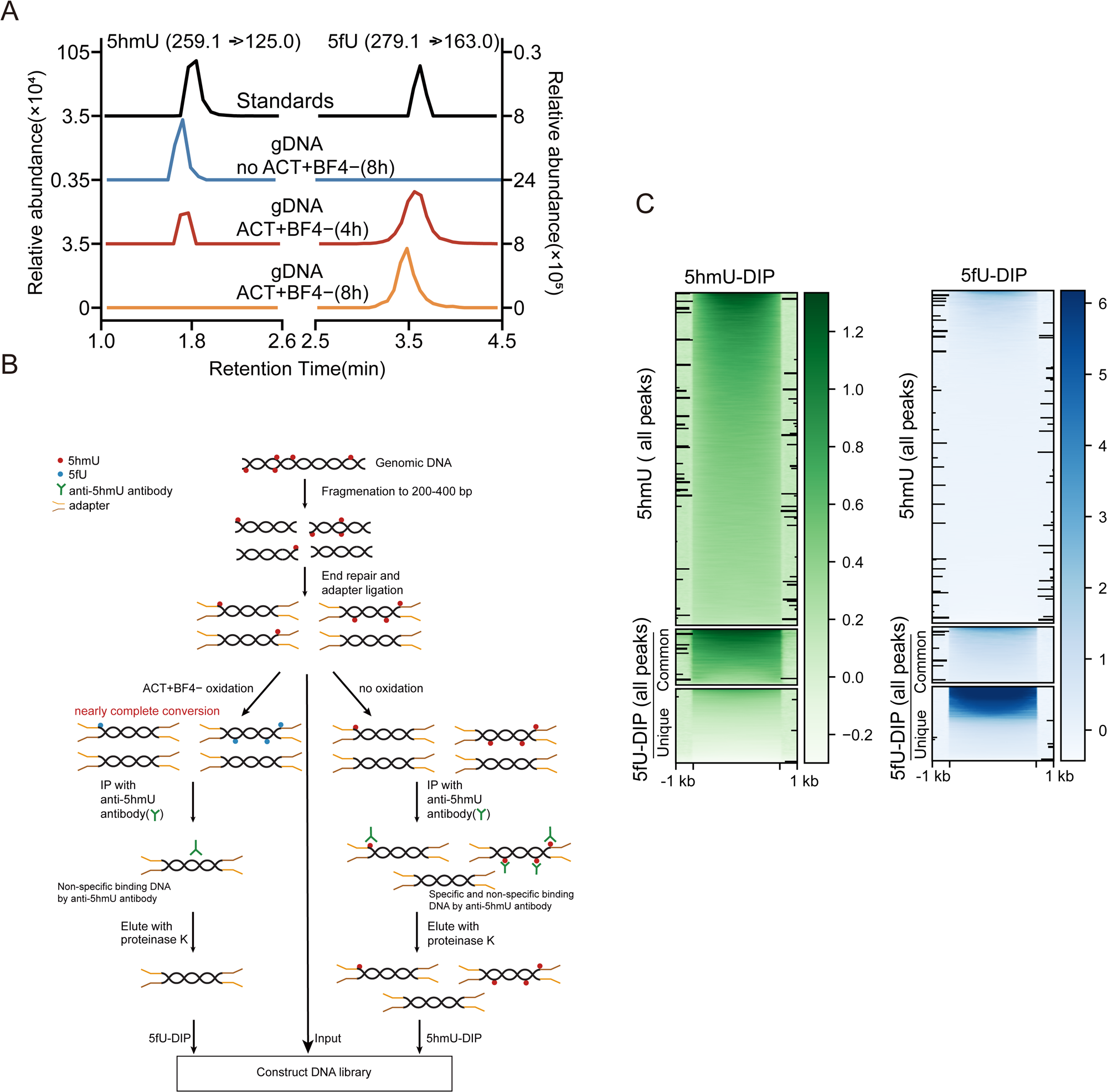
DNA immunoprecipitation analysis of 5hmU in the *A. carterae* genome. (A) LC-MS/MS chromatograms showing the complete conversion of 5hmU to 5fU at the dsDNA level after ACT^+^BF4^−^ treatment. (B) Schematic diagram of 5hmU-DIP and 5fU-DIP. (C) Heatmap of signal intensities for 5hmU-DIP and 5fU-DIP peaks with 1-kb upstream and downstream flanks identified in *A. carterae* genome. Common and Unique, represent the 5fU-DIP peaks which shared overlapped or non-overlapped positions with 5hmU-DIP peaks respectively.

**Figure S6.**
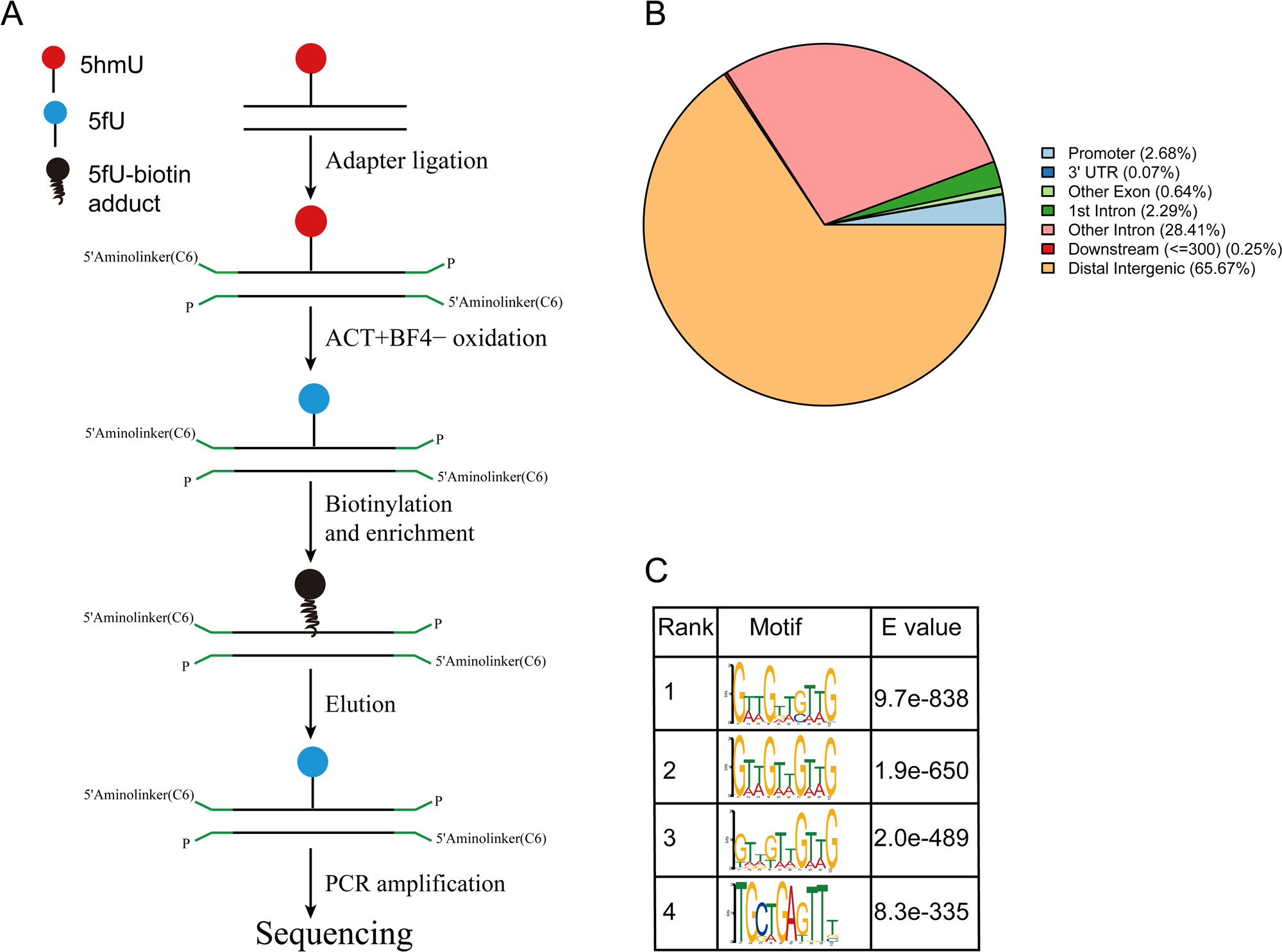
Global profile of 5hmU peaks in *A. carterae* genome obtained by chemical pull-down approach. (A) Schematic diagram of the chemical-tagging strategy for genome-wide 5hmU loci enrichment in *A. carterae*. (B) Distribution of 5hmU peaks across different genomic regions in the *A. carterae* genome. (C) Sequence motifs enriched at 5hmU-specific loci. The top four motifs identified are shown with their respective E values.

**Figure S7.**
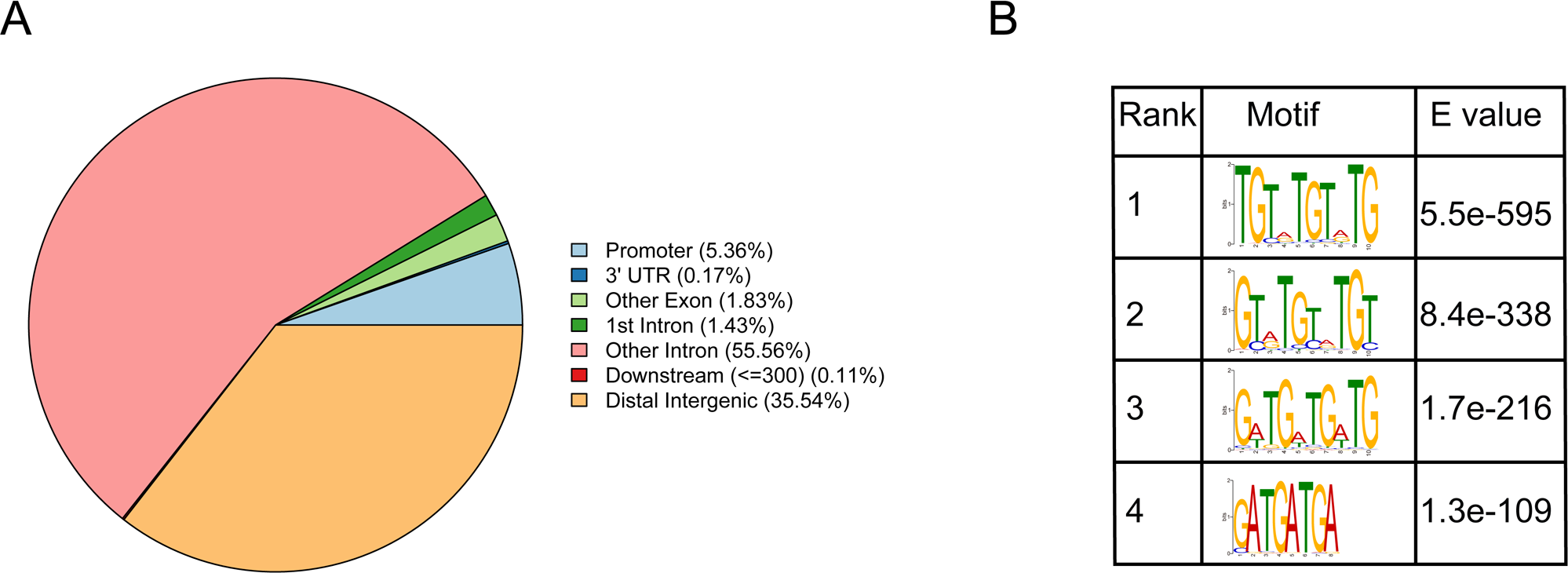
The 5hmU profile and its conserved motifs in the *Symbiodinium* sp. genome. (A) The genomic distribution of 5hmU peaks across different genomic regions in *Symbiodinium* sp.. (B) The conserved motifs present in the enriched 5hmU loci.

**Figure S8.**
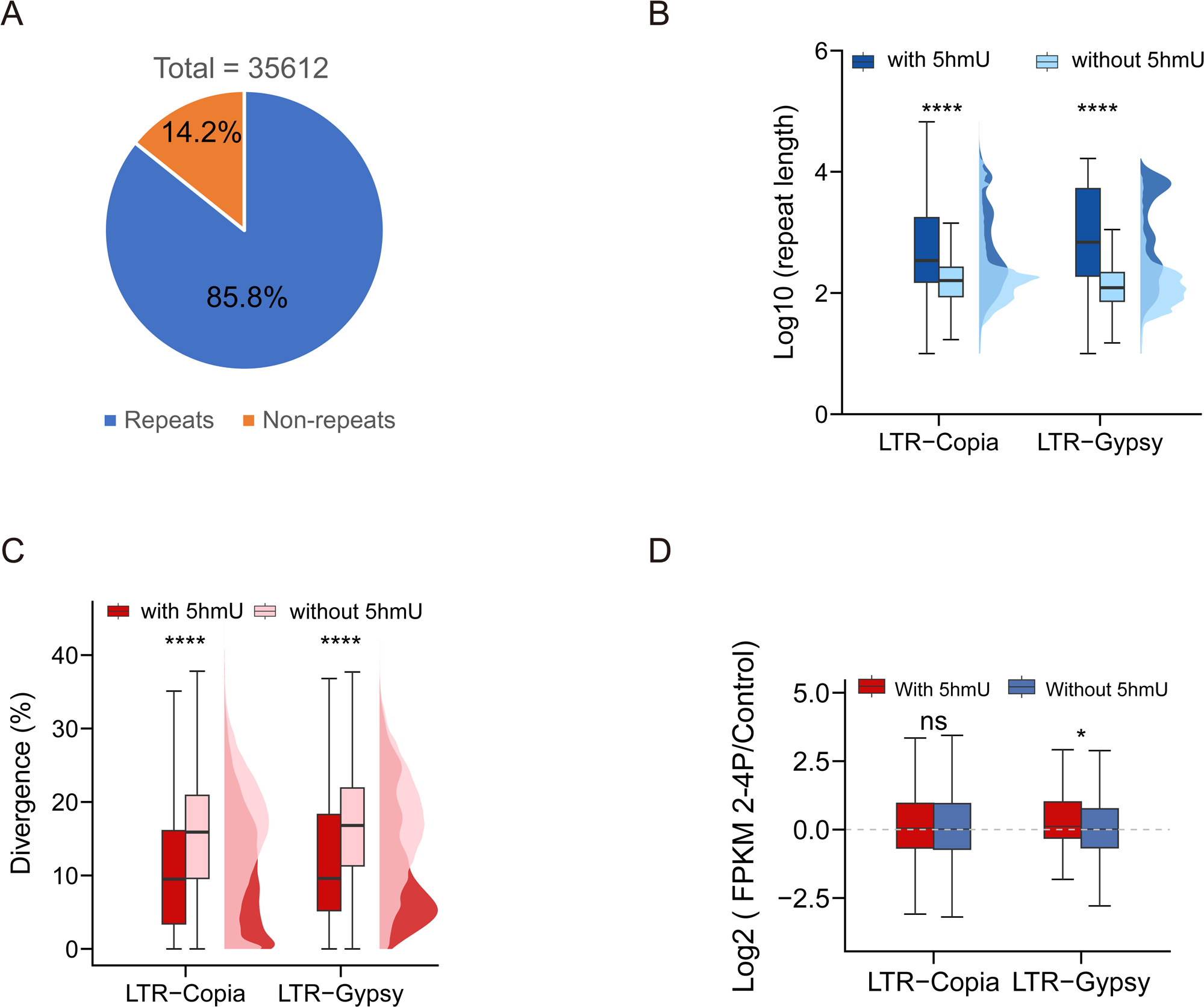
The 5hmU distribution in *A. carterae* genome-wide repeats, and the features of LTR elements containing with or without 5hmU peaks. (A) Pie chart showing percentages of 5hmU peaks over repeat and non-repeat sequences in *A. carterae* genome. (B and C) The length (B), and corresponding divergence (C) of LTR-Copia (Copia) and LTR-Gypsy (Gypsy) with or without 5hmU peaks (Wilcox test, ****p < 0.0001). (D) The Log2 fold change of transcriptional expression levels (FPKM values) for *A. carterae* cells after 2-4P treatment regarding both Copia and Gypsy elements with or without 5hmU peaks (Wilcox test, ns = not significant; *p < 0.05). Control, *A. carterae* cells with no 2-4P treatment. 2-4P, *A. carterae* cells after 2-4P treatment. For all boxplots, the center lines represent median values. The box limits denote upper (third) and lower (first) quartiles, while the error bars represent the highest and lowest values.

**Figure S9.**
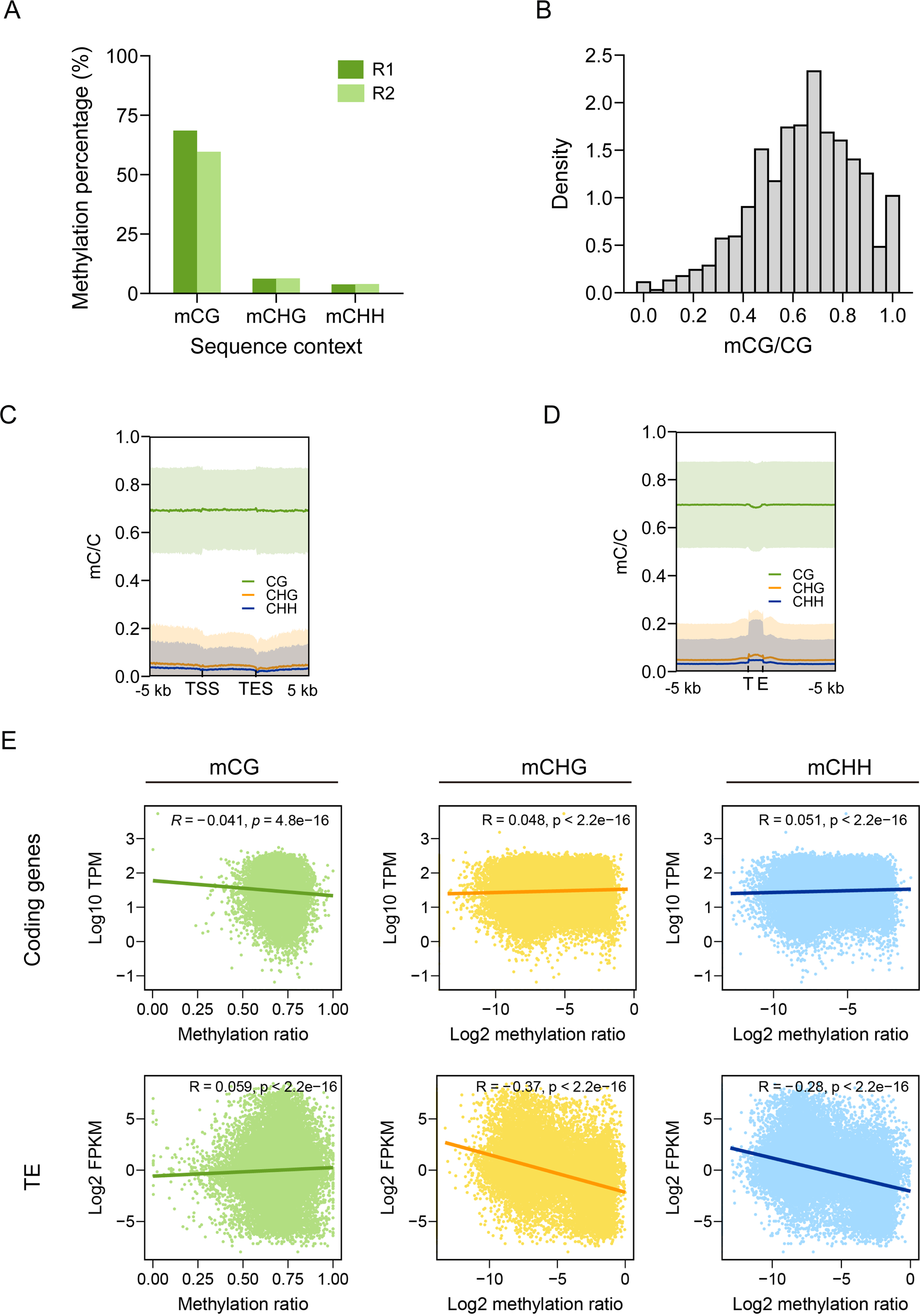
Characteristics of genome-wide 5mC patterns in dinoflagellate *A. carterae*. (A) The cytosine methylation level in different sequence contexts including CG, CHG and CHH. R1 and R2 represent two biological samples. (B) The mCG density histogram of individual CG sites with high coverage (≥10 reads). (C and D) Cytosine methylation profiles at CG, CHG, and CHH sites on *A. carterae* genes and transposable elements (TEs). Thick lines indicate mean values and pale shades show standard deviation. TSS and TES are abbreviations of transcriptional start site and transcriptional end site respectively. (E) Correlation analyses of cytosine methylation level with transcriptional levels of genes and TEs.

**Figure S10.**
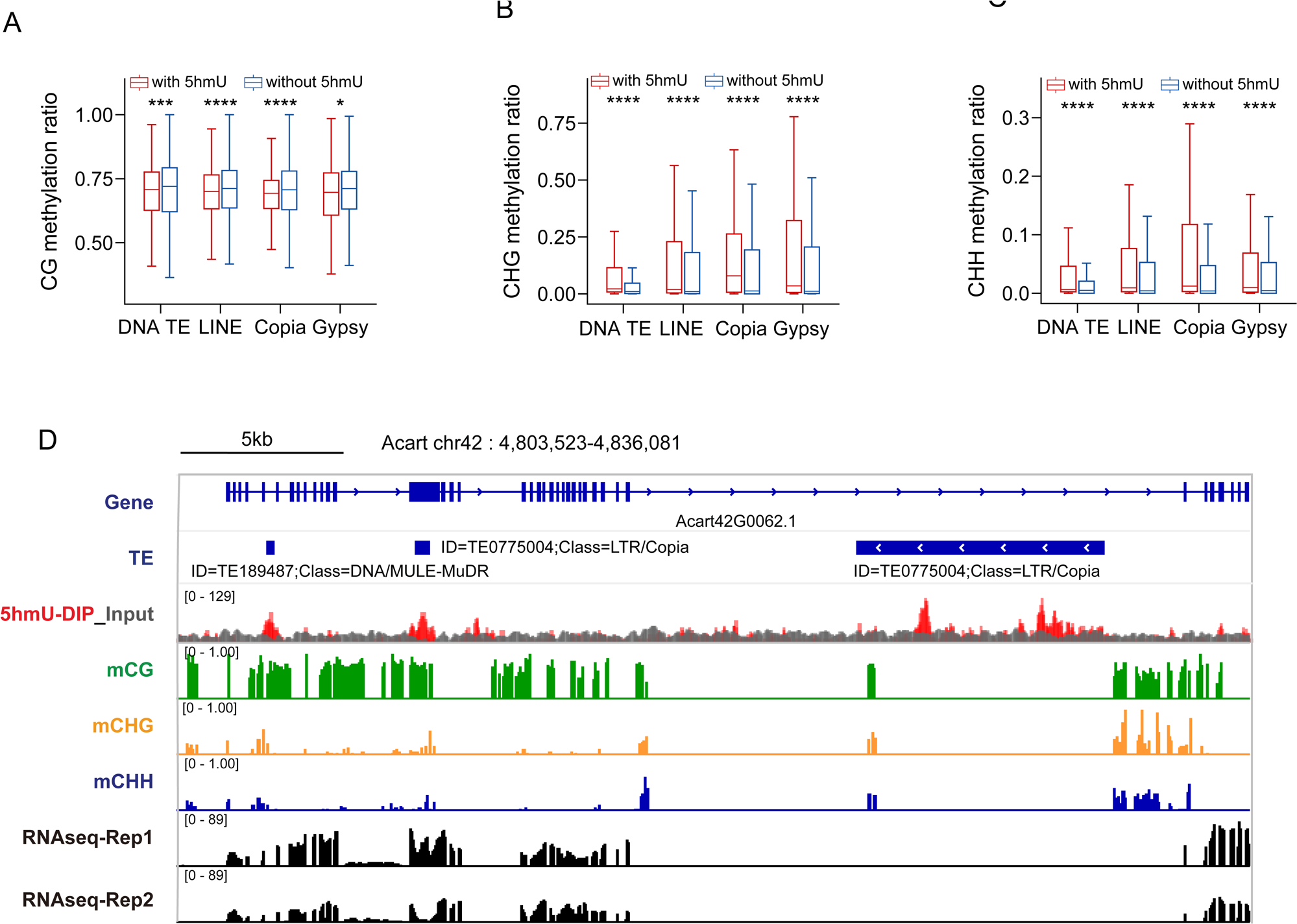
5mC levels in different types of transposable elements bearing with or without 5hmU peaks. (A-C) The DNA methylation ratios of mCG (A), mCHG (B), mCHH (C) in repeat elements including DNA TE, LINE, Copia and Gypsy (Wilcox test, *p <0.05; ***p <0.001; ****p <0.0001). For all boxplots, the centerlines indicate medians, box edges represent the interquartile range and the outliers are not shown. (D) Genome browser showing 5hmU loci and 5mC deposition in a region of *A. carterae* genome. RNAseq-Rep1 and RNAseq-Rep1 represent RNAseq data from two biological replicates of *A. carterae* cells treated with no 2-4P.

**Figure S11.**
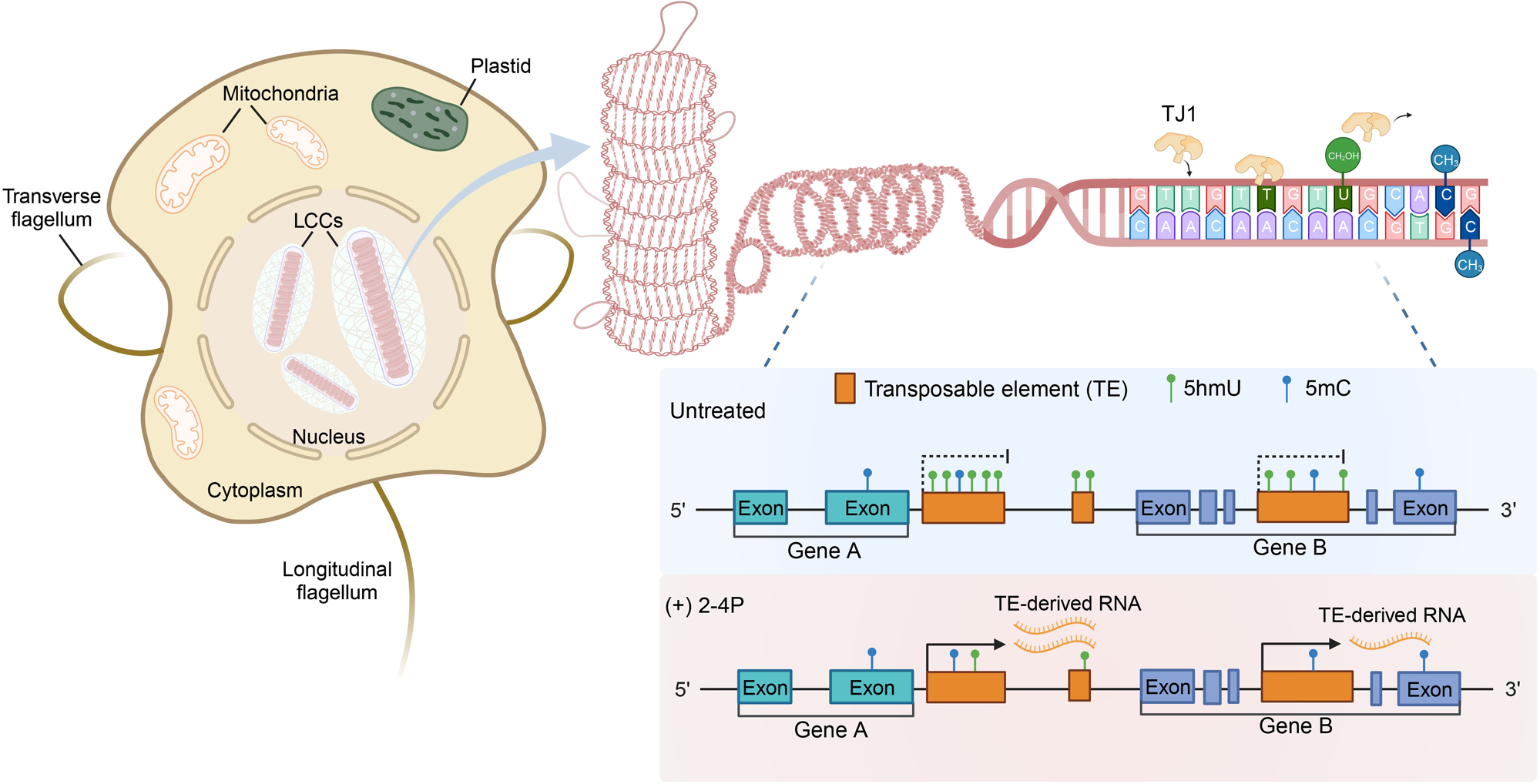
Hypothetical model showing the source and putative function of 5hmU in dinoflagellates. With preferential occurrence within sequence context of GTTGTTGTTG, the unusually high amounts of 5hmU in dinoflagellate LCCs, possibly derived from dinoflagellate TET/JBP homologues TJ1 proteins-mediated oxidation of thymidine at the polynucleotide level, are primarily distributed in intergenic and intronic regions, and also showed main co-colocalization with transposable elements (TEs). Additionally, the decrease of global 5hmU content through using 5hmU synthase inhibitor (2-4P) leads to the activation of the 5hmU-marked TEs, implicating this unusual DNA modification may be used as epigenetic mark to silence transposons in dinoflagellates.

